# Whole blood transcriptional profiles and the pathogenesis of tuberculous meningitis

**DOI:** 10.1101/2023.10.06.561265

**Authors:** Hoang Thanh Hai, Le Thanh Hoang Nhat, Trinh Thi Bich Tram, Do Dinh Vinh, Artika P Nath, Joseph Donovan, Nguyen Thi Anh Thu, Dang Van Thanh, Nguyen Duc Bang, Dang Thi Minh Ha, Nguyen Hoan Phu, Ho Dang Trung Nghia, Le Hong Van, Michael Inouye, Guy E Thwaites, Nguyen Thuy Thuong Thuong

## Abstract

**Background:** Mortality and morbidity from tuberculous meningitis (TBM) are frequent and strongly associated with the inflammatory response to *Mycobacterium tuberculosis* infection. However, the mechanisms driving the associations are uncertain. We sought to identify the gene modules, hubs and pathways associated with the pathogenesis and mortality from TBM, and to identify which best-predicted death.

**Methods:** We used whole blood RNA sequencing to obtain transcriptional profiles from 281 Vietnamese adults with TBM (207 HIV-negative; 74 HIV-positive), 295 with pulmonary TB (PTB), and 30 healthy controls. The TBM cohort was divided randomly into a discovery cohort (n=142) and a validation cohort (n=139). Weighted gene co-expression network analysis identified clusters of genes (or ‘modules’) and hub genes associated with death or disease severity. An overrepresentation analysis identified pathways associated with TBM mortality, with a consensus analysis identifying consensual patterns between HIV-positive and HIV-negative individuals. A multivariate elastic-net Cox regression model selected the candidate predictors of TBM mortality, then model prediction performance using logistic regression and internal bootstrap validation to choose best predictors.

**Results:** Overall, TBM mortality was associated with increased neutrophil activation and decreased T and B cell activation pathways. Death from TBM was associated with increased angiogenesis in HIV-positive adults, and with activated TNF signaling and down-regulated extracellular matrix organization in HIV-negative adults. PTB and TBM have similar transcriptional profiles compared to healthy controls, although inflammatory genes were more activated in HIV-positive than HIV-negative TBM. The expression of four hub genes – *MCEMP1*, *NELL2*, *ZNF354C* and *CD4* – were strongly predictive of death from TBM (AUC 0.80 and 0.86 for HIV-negative and HIV-positive, respectively).

**Conclusions:** Whole blood transcriptional profiling revealed that TBM is associated with a characteristic systemic inflammatory response, similar to that invoked by pulmonary tuberculosis, but with key gene modules, hubs and pathways strongly associated with death. Our analysis suggests a novel 4-gene biomarker for predicting death from TBM, but also opens a new window into TBM pathogenesis that may reveal novel therapeutic targets for this lethal disease.

## Introduction

Of 7.1 million new tuberculosis (TB) cases in 2019, tuberculous meningitis (TBM) is estimated to have developed in 164,000 adults, around 25% of whom were living with HIV ^1^. TBM is the most severe form of TB, causing death or neurological disability in half of all cases. Overall, TBM mortality is around 25%, but rises to around 50% in those with HIV, with most deaths occurring in the first 3 months of treatment ^1,2^.

The poor outcomes from TBM are strongly associated with the inflammatory response ^3,4^, with both a paucity and an excess of inflammation linked to death from TBM ^5–11^. However, the mechanisms behind these observations remain uncertain. Immune responses in TBM are thought to be compartmentalized within the central nervous system. Studies have shown that immune cell counts, cytokine concentrations, metabolites and transcriptional responses differ between the peripheral blood and the cerebrospinal fluid (CSF) ^10,12,13^. In adults with TBM, leukocyte activation is higher in the CSF than in peripheral blood, although a marked myeloid response in peripheral blood has been reported ^10^. Blood transcriptomic analysis has found increased neutrophil-associated transcripts and inflammasome signaling in those with HIV-associated TBM and immune reconstitution inflammatory syndrome ^5^. In children with TBM, whole blood transcriptional profiles showed increased inflammasome activation and decreased T-cell activation ^12^. Taken together, these studies suggest the inflammatory response associated with TBM may have a greater systemic component than originally thought and its characterization may help define the immune mechanisms leading to death and disability.

The accurate and early detection of patients at highest risk of complications and death from TBM may help to target treatment for those most in need. The British Medical Research Council (MRC) grades have been used to categorize TBM severity for almost 80 years ^14^, and the system strongly predicts TBM mortality ^15^. Previously, we and others developed new prognostic models from studies of 1699 adults with TBM using clinical and laboratory parameters, including MRC grade ^4,15^. These models predicted outcome more accurately than MRC grade alone. However, they might be improved by measures of host inflammatory response. Host-based peripheral blood gene expression analysis has been used to identify active or progressive pulmonary TB and in pulmonary TB treatment monitoring ^16,17^, but has yet to be applied to TBM.

In the current study, we investigated whole blood RNA sequencing (RNA-seq) transcriptional profiles in 281 Vietnamese adults with TBM, 295 with pulmonary TB (PTB), and 30 healthy controls. Our objective was to use weighted gene co-expression network analysis, an unbiased and well-evaluated approach, to identify the biological pathways and hub genes associated with TBM pathogenesis and assess the predictive value of gene expression for early mortality from TBM.

## Results

### Characteristics and outcomes of the cohorts

Four RNA-seq cohorts (all ≥18 years) were used in the study, representing a total of 606 participants. The characteristics of these cohorts are provided in **Table 1**. There were 281 adults with TBM; 207 HIV-negative and 74 HIV-positive. In the HIV-negative TBM adults, the median age was 46 years (IQR 34, 58), 127 (61%) were male, and the median Body Mass Index (BMI) was 20.0 (IQR 18.2, 22.3). HIV-positive TBM were more likely than HIV-negative TBM to be male, younger, have lower BMI, have previously received TB treatment, and to have microbiologically confirmed TBM. Total white cell counts in blood and CSF in HIV-positive TBM were lower than in HIV-negative TBM. Median CD4 cell counts in HIV-positive TBM were 67 cells/mm^3^ (IQR 19, 124) and 28 (39%) were receiving antiretroviral therapy. The PTB cohort consisted of 295 HIV-negative adults with the median age of 44 years (IQR 31, 52), 228 (77%) were male, the median of BMI was 19.4 (IQR 17.7, 21.6) and 129 (48%) had pulmonary cavities on chest X-ray. Of the 30 healthy controls, 11 (37%) were male, and the median age was 33 (IQR 29, 37). In real-time quantitative polymerase chain reaction (qPCR) validation cohort, 132 HIV-negative TBM adults have similar characteristics as HIV-negative TBM RNA-seq cohort (**Table 1**).

**Table 1.** Baseline characteristics of TBM, PTB, and healthy controls.

| Characteristics | RNA-seq cohorts |  |  |  |  |  |  |  | qPCR validation cohort |  |
| --- | --- | --- | --- | --- | --- | --- | --- | --- | --- | --- |
|  | HIV-negative TBM<br>n=207 |  | HIV-positive TBM<br>n=74 |  | PTB<br>n=295 |  | Healthy controls<br>n=30 |  | HIV-negative TBM<br>n=132 |  |
|  | n | Summary | n | Summary | n | Summary | n | Summary | n | Summary |
| **Age (years)** | 207 | 46 (34, 58) | 74 | 34 (29, 40) | 295 | 44 (31, 52) | 30 | 33 (29, 37) | 132 | 48 (35, 60) |
| **Male sex** | 207 | 127 (61) | 74 | 56 (76) | 295 | 228 (77) | 30 | 11 (37) | 132 | 84 (65) |
| **BMI (kg/m<sup>2</sup>)** | 205 | 20.0 (18.2, 22.3) | 72 | 19.3 (17.2, 20.4) | 295 | 19.4 (17.7, 21.6) |  |  | 132 | 20.0 (18.2, 22.3) |
| **Symptom duration (days)** | 207 | 14 (11, 20) | 73 | 16 (10, 30) | 294 | 20 (10, 30) |  |  | 132 | 16 (13, 24) |
| **History of TB treatment** | 204 | 5 (2.5) | 74 | 15 (20) | 295 | 100 (34) |  |  | 130 | 12 (9.2) |
| **Glasgow coma score** | 207 | 14 (12, 15) | 72 | 14 (13, 15) |  |  |  |  | 132 | 14 (13, 15) |
| **Cavity chest X-ray** |  |  |  |  | 270 | 129 (48) |  |  |  |  |
| **TB microbiological tests** |  |  |  |  |  |  |  |  |  |  |
| MGIT culture positive | 199 | 50 (25) | 69 | 41 (59) | 295 | 279 (95) |  |  |  |  |
| Xpert/Ultra positive | 198 | 42 (21) | 70 | 39 (56) | 295 | 287 (97) |  |  |  |  |
| Microscopy positive | 205 | 48 (23) | 67 | 35 (52) | 203 | 169 (83) |  |  | 105 | 18 (17) |
| **Blood (10<sup>6</sup> cells/ml)** |  |  |  |  |  |  |  |  |  |  |
| Leucocyte count | 204 | 9.4 (7.0, 11.9) | 74 | 6.4 (5.0, 9.2) | 242 | 9.2 (7.4, 11.4) | 26 | 6.4 (5.6, 7.2) | 129 | 10.0 (7.7, 12.4) |
| Neutrophil count | 204 | 7.1 (4.8, 9.1) | 74 | 5.0 (3.3, 6.9) | 241 | 6.1 (4.7, 8.2) | 26 | 3.4 (3.1, 4.1) | 129 | 7.8 (6.9, 8.5) |
| Lymphocytes count | 204 | 1.2 (0.9, 2.0) | 74 | 0.7 (0.4, 1.2) | 242 | 1.9 (1.4, 2.3) | 26 | 2.2 (1.9, 2.6) | 129 | 1.2 (0.7, 1.8) |
| **CSF (10<sup>3</sup> cells/ml)** |  |  |  |  |  |  |  |  |  |  |
| Leucocyte count | 207 | 142 (19, 323) | 73 | 124 (10, 453) |  |  |  |  | 106 | 122 (38, 328) |
| Neutrophil count | 207 | 0 (0, 39) | 73 | 17 (0, 144) |  |  |  |  | 60 | 20 (3, 73) |
| Lymphocyte count | 207 | 106 (18, 223) | 73 | 58 (10, 216) |  |  |  |  | 106 | 95 (79, 100) |
| **CD4 cell count (cells/ml)** |  |  | 73 | 67 (19, 124) |  |  |  |  |  |  |
| **Antiretroviral therapy** |  |  | 70 | 28 (39%) |  |  |  |  |  |  |
| **HIV load (10<sup>3</sup> cells/ml)** |  |  | 70 | 77.3 (8.28, 672) |  |  |  |  |  |  |

Values were displayed as median (1<sup>st</sup> and 3<sup>rd</sup> interquartile) for continuous variables and frequency (%) for categorical variables.
TBM = tuberculous meningitis, PTB= pulmonary tuberculosis, CSF= cerebrospinal fluid

The clinical variables associated with three-month mortality of the 281 adults with TBM in RNA-seq cohort are given in **Table 2**. The discovery (n=142) and validation (n=139) cohorts had similar characteristics (**Table 2**). 47.5% (133/280) had definite TBM ^18^, with microbiologically confirmed disease, accounting for 45.9% (101/220) of survivors and 52.4% (32/61) of those who died. The overall three-month mortality rate was 21.7% (61/281) for TBM regardless of HIV status: 16.4% (34/207) in HIV-negative and 36.5% (27/74) in HIV-positive (p<0.001). We did not observe differences in mortality by sex, age and diagnostic category. Greater disease severity, MRC grades 2 and 3 at enrolment, was associated with increased mortality compared to grade 1 (p<0.001). In those who died, CSF and peripheral blood neutrophil counts were higher and peripheral blood lymphocyte count lower, compared to those who survived.

**Table 2.**
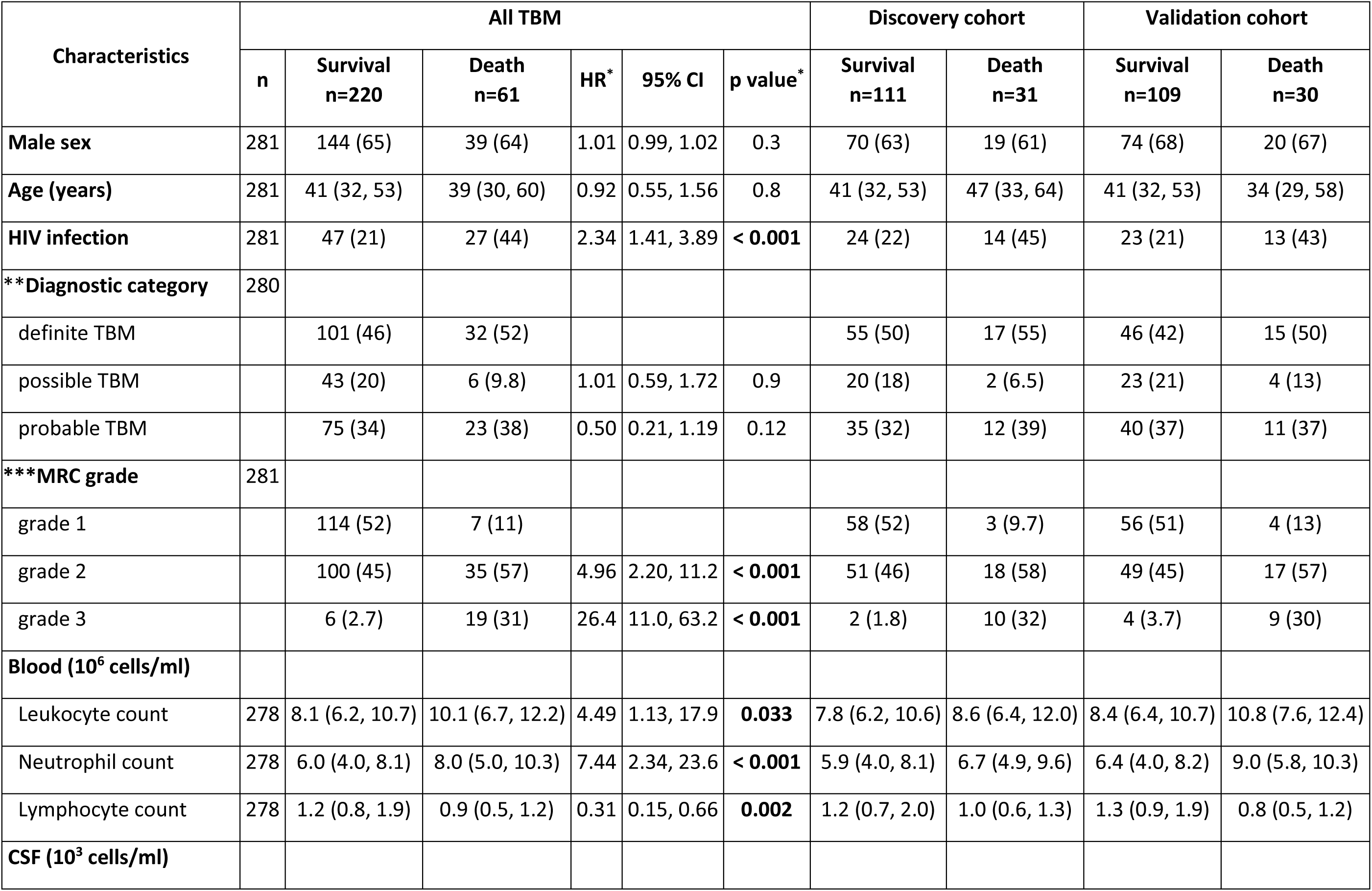
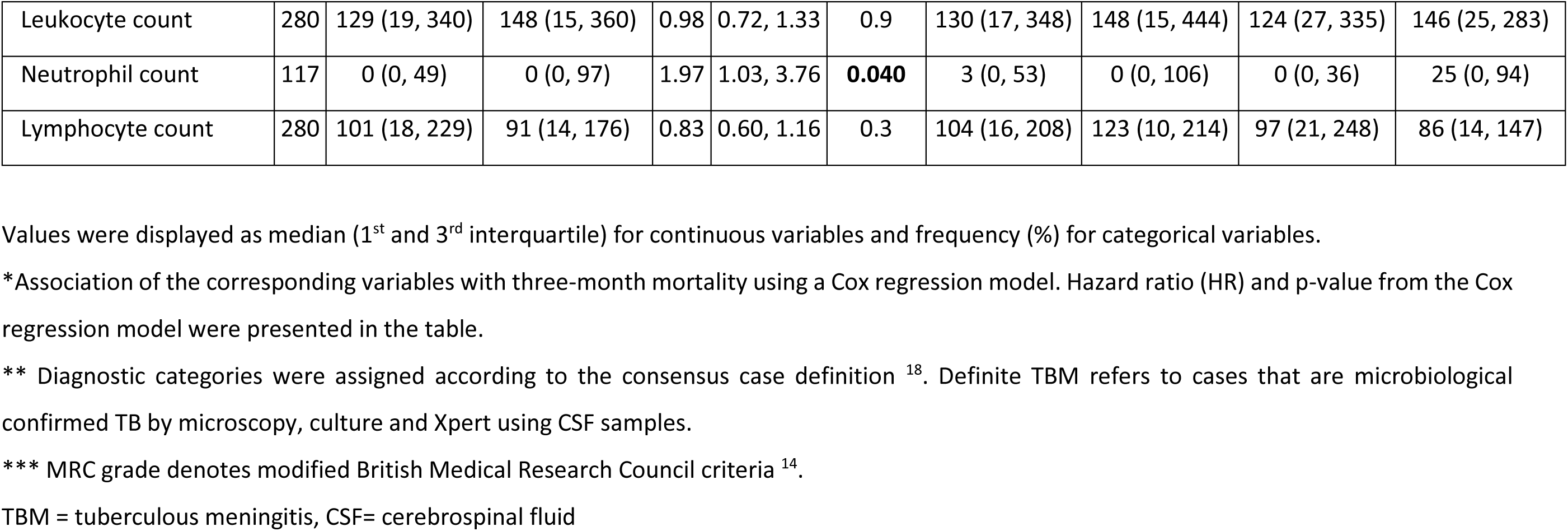
Association between baseline clinical characteristics with TBM mortality in RNA-seq cohorts.

In the qPCR validation cohort (n=132), three-month mortality rate was 28.8% (38/132) and those who died were of older age, had more severe disease, and had lower number of CSF leukocytes and lymphocytes (Table S1). Death was not associated with CSF or peripheral blood neutrophil counts.

### Whole blood transcriptional profiles of the four RNA-seq cohorts

We analyzed the whole blood transcriptomics, using bulk RNA sequencing from 606 participants in the 4 cohorts. On average 35.1 million reads/sample were obtained with 89.4% reads mapping accuracy to human reference genome (GRCh.38 release 99) and 65.4% of reads were uniquely mapped. The study objectives and cohorts flow are presented in **Figure 1**. Principal component analysis on the transcriptomic data of 20,000 genes across 4 studies showed different profiles between healthy controls from both PTB and TBM cases (**Figure 2**). The PTB profile substantively overlapped with TBM, with some separation between HIV-negative and HIV-positive TBM (**Figure 2**).

**Figure 1.**
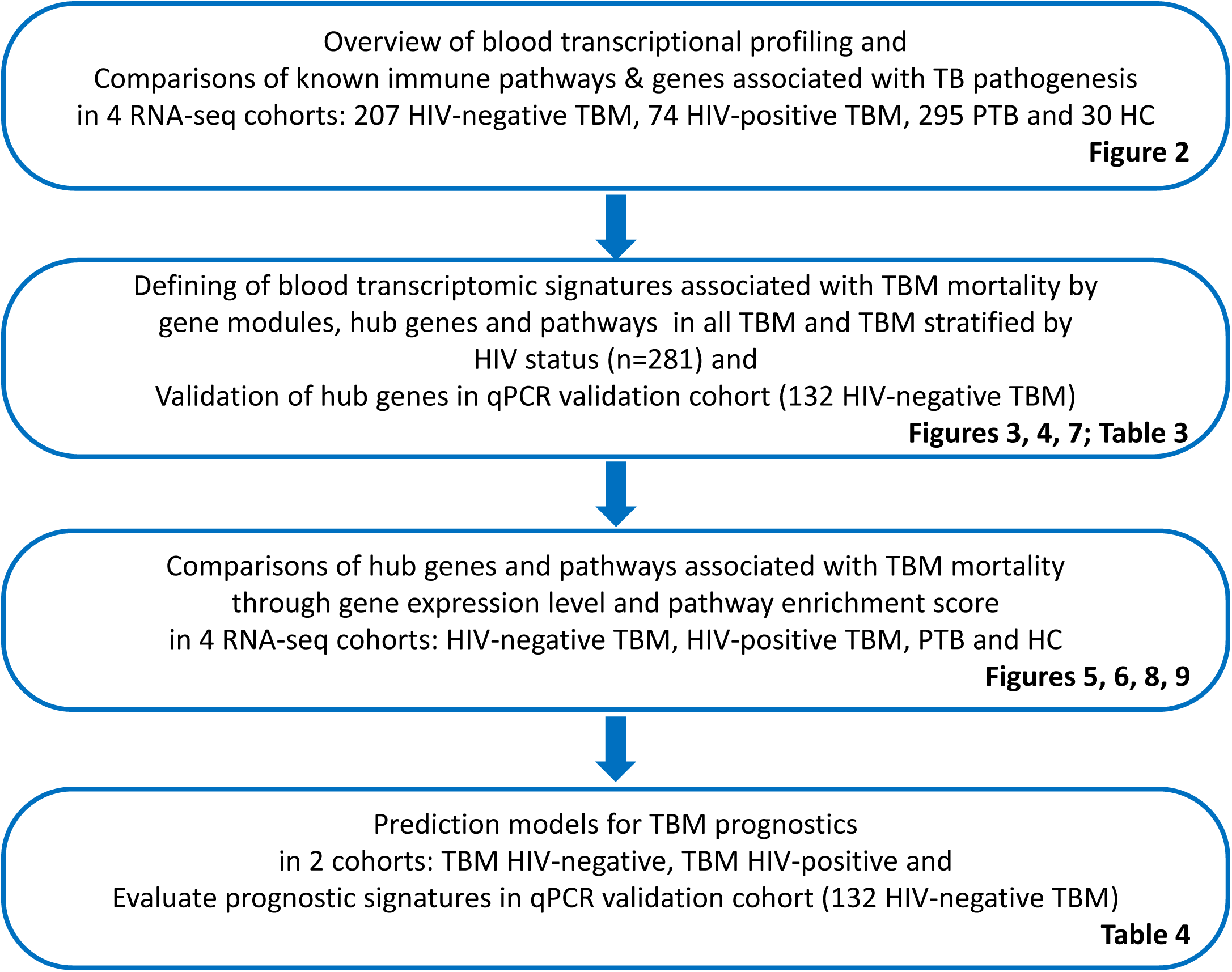
Objectives and cohorts flow. TBM: TB meningitis, HIV: human immunodeficiency virus, PTB: pulmonary TB, HC; healthy controls

**Figure 2:**
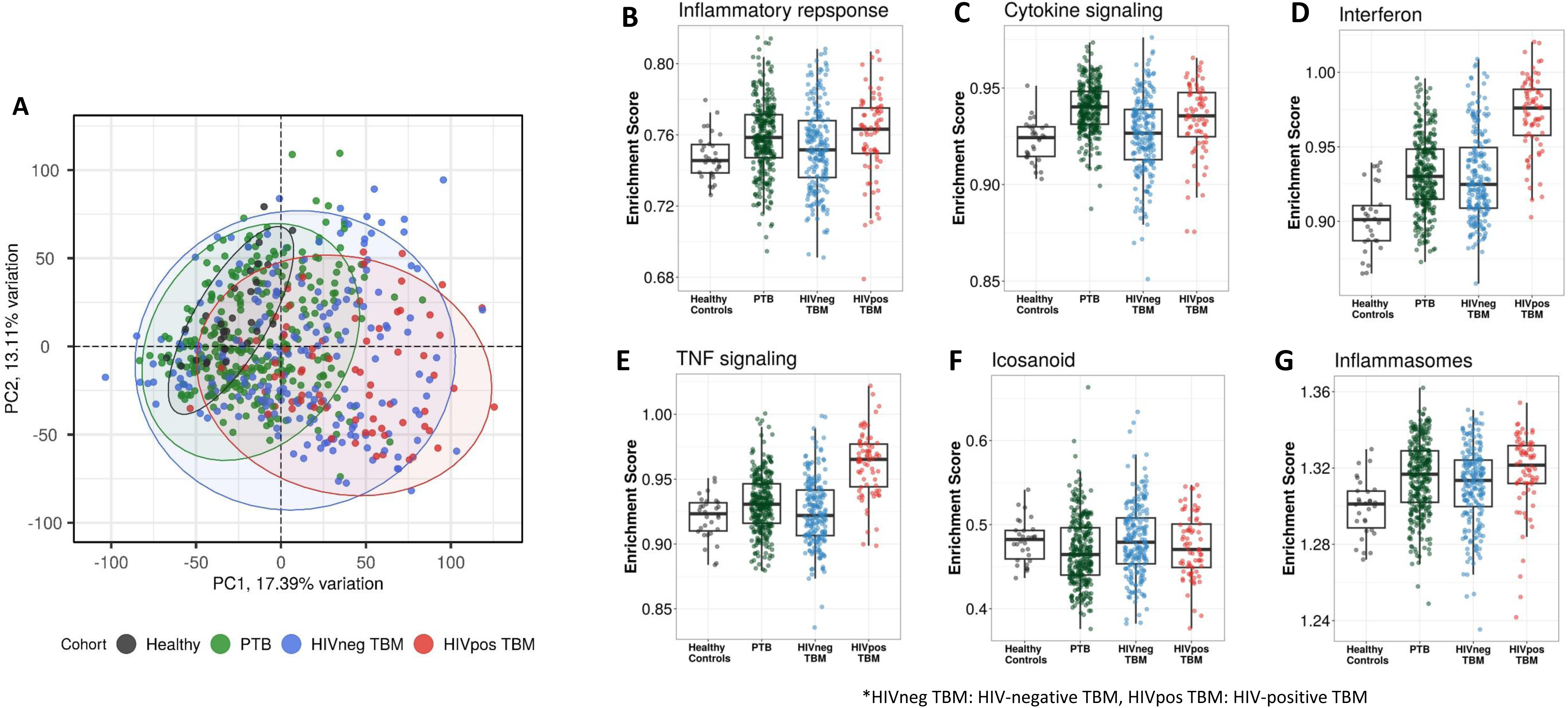
Blood transcriptomic profiles of four cohorts: healthy controls (n=30), PTB (n=295), HIV-negative TBM (n=207) and HIV-positive TBM (n=74) **(A)** Principle component analysis (PCA) of whole transcriptomic profile of HC, PTB PTB and TBM with and without HIV. Each symbol represents one individual with color coding different cohorts. The x-axis represents principle component (PC) 1, while y-axis represents PC2. **(B-G)** Enrichment scores of known innate immunity pathways associated with TBM pathogenesis. Gene list of these pathway were depicted in additional file 1, table S5. Pathway enrichment scores were calculated using single sample GSEA algorithm (ssGSEA) (Barbie DA, et al 2009). Each dot represents one participant. The box presents median, 25^th^ to 75^th^ percentile and the whiskers present the minimum to the maximum points in the data.

Enrichment scores from single sample gene set enrichment analysis (ssGSEA), which based on expression rank of genes relevant to pathways, were measured in some pathways already known to be important mediators of TB or TBM pathogenesis (**Figure 2**). As anticipated, inflammatory response, cytokine signaling, interferon signaling, TNF signaling, and inflammasome activation pathways, were enriched in PTB and TBM cohorts as compared to healthy controls. In TBM, enrichment of genes in these pathways were generally higher in HIV-positive than in HIV-negative individuals (**Tables S2, S6**).

### Transcriptional gene modules associated with TBM severity and mortality

Transcriptional profiles associated with TBM mortality were generated by identifying differentially expressed genes. Of the top 20,000 genes with most variation, we observed 724 (3.6%) genes that were differentially expressed (FDR <0.05, absolute fold change (FC) >1.5) in all those with TBM (**Figure 3, Table S3**). Next, we applied weighted gene co-expression network analysis (WGCNA) to 5000 most variable genes from 281 TBM samples (n= 207, HIV-negative; n=74, HIV-positive) to define clusters of highly correlated genes (modules) associated with TBM severity and mortality. Gene modules are clusters of genes that have highly interconnected expression, usually related to their biological functions. Hub genes are genes with high connectivity to other genes within a respective module. First, we used WGCNA to construct a network of gene modules in the discovery cohort. Then we validated the presence of these transcriptional modules in the validation cohort, labelling the modules with different colors. In the discovery cohort (n=142), 15 modules were identified overall, consisting of 44 to 1350 genes per module (**Figure S1, Table S7**). All 15 modules were preserved in the validation cohort (n=139) through the preservation analysis (**Figure S2**).

**Figure 3:**
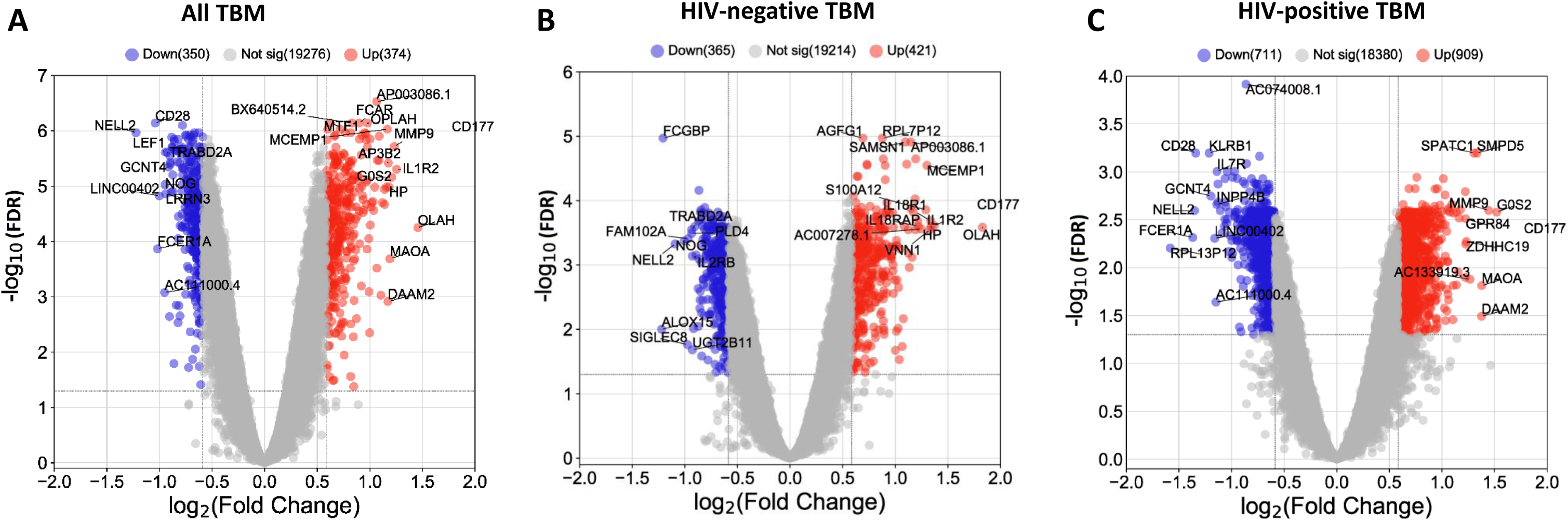
Blood transcriptomic profiles of three-month mortality at baseline in all TBM and TBM stratified by HIV status. Volcano plot showed differential expression (DE) genes by fold change (FC) between death and survival in all TBM **(A)**, HIV-negative **(B)** and HIV-positive TBM **(C)**. Each dot represents one gene. The x-axis represents log_2_ FC. The y-axis showed –log_10_ FDR of genes. DE genes were colored with red indicating up-regulated, blue indicating down-regulated genes which having fold discovery rate (FDR) <0.05 and absolute FC > 1.5.

The associations between the 15 modules and TBM severity and mortality are presented in **Figure 4** for both the discovery and validation cohorts. Modules were linked to each other in a hierarchical structure, with major biological processes annotated. Associations of the modules with TBM disease severity (MRC grade) at baseline were estimated by Spearman correlations between MRC grade and the first principle component (PC1) of each module. Similarly, associations of the modules with mortality were measured by hazard ratio (HR) per increase 1 unit of PC1 using Cox regression model adjusted for age, HIV status and dexamethasone treatment (**Figure 4**) in both discovery and validation cohorts.

**Figure 4.**
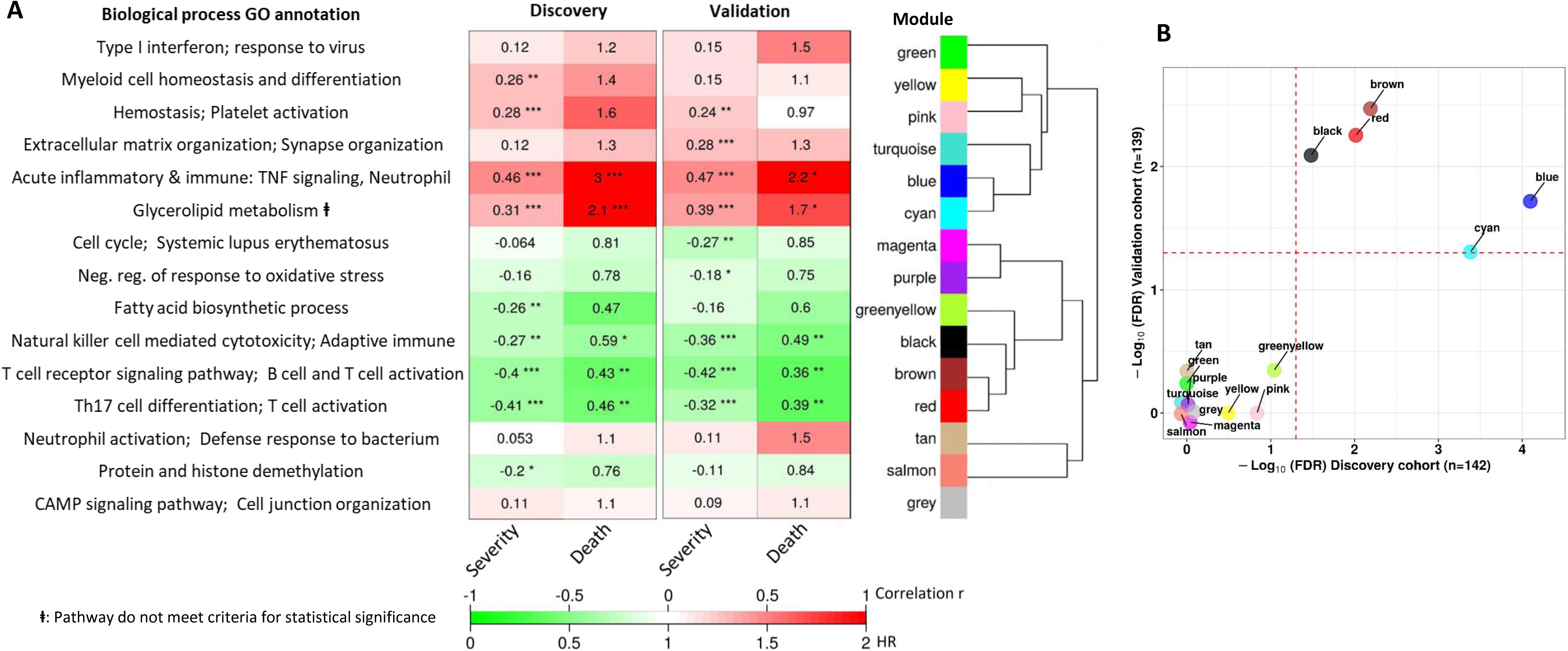
Blood transcriptional modules associated with mortality in TBM. **(A)** Associations between WGCNA modules with two clinical phenotypes TBM disease severity (MRC grade) and three-month mortality in discovery and validation cohorts, and their associated biological processes. The heatmap showed the association between principle component 1 (PC1) of each module and the phenotypes, particularly Spearman correlation r for MRC grade and hazard ratio per increase 1/10 unit of PC1 (HR) for mortality. The HRs were estimated using a Cox regression model adjusted for age, HIV status and dexamethasone treatment. False discovery rate (FDR) corrected based on Benjamini & Yekutieli procedure, with significant level denoted as * < .05, ** < .01 and *** <.001. Gradient colors were used to fill the cell with green indicating negative r or HR < 1, red color indicating positive r or HR > 1. The order of modules was based on hierarchical clustering using Pearson correlation distance for module eigengene. On the left, biological processes, corresponding to modules, were identified using Gene Ontology and KEGG database. **(B)** Validation of the association between WGCNA modules and mortality in discovery and validation cohorts. X-axis represents –log_10_ FDR in discovery cohort and Y-axis represents –log_10_ FDR in validation cohort. Red dash lines indicate FDR = 0.05 as the threshold for statistically significant in both cohorts. Five modules (blue, brown, red, black and cyan) with FDR < 0.05 were validated.

Of the 15 preserved modules, five modules were significantly associated with mortality in the discovery and validation cohorts, with false discovery rate (FDR) < 0.05 (**Figure 4**). These five modules were separated into two big module clusters. The first cluster contained the blue module (n=799 genes), involved in inflammatory and innate immune responses, and the cyan module (n=44 genes) with unknown biological function. These modules were upregulated in those who died, as shown in the heat-map in **Figure 4** (HR: 3.0 and 2.2 for the blue module, and 2.1 and 1.7 for the cyan module, FDR < 0.05 for all comparisons). The black (n= 207 genes), brown (n=698 genes) and red (n=229 genes) modules were in the second cluster and were generally involved in adaptive immunity including T and B cell signaling pathways. These three modules were down-regulated in those who died (HR: 0.43 and 0.36 for brown, 0.46 and 0.39 for red, 0.59 and 0.49 for black) (**Figure 4**).

It is known that TBM MRC grade before treatment initiation strongly predicts outcome from TBM. Here, we investigated correlations between each module and MRC grade and their association with mortality. The pink module, involved in hemostasis and platelet activation, was positively correlated with MRC grade, but not mortality. Of the five modules associated with death from TBM, all were correlated with MRC grade (**Figure 4**). Four of these five modules, were enriched for immune responses.

### Transcriptional hub genes associated with TBM severity and mortality

We next identified hub genes within the four biologically functional modules associated with TBM mortality in both the discovery and validation cohorts. Hub genes showed higher connectivity within the modules, and stronger association with TBM mortality as compared to less connected genes within a module (**Figure S3**). Seven hub genes (*ETS2, PGD, UPP1, CYSTM1, FCAR, KIF1B* and *MCEMP1*) were upregulated in death, all from the acute inflammation (blue) module. Hub genes from the brown, red and black modules were downregulated in death and were involved in adaptive immune response. Ten hub genes associated with mortality (*CD96, TNFRSF25, TBC1D4, CD28, ABLIM1, RASGRP1, NELL2, TRAF5, TESPA1, TRABD2A*) were from the brown module, three (*EVL, PLCG1, NLRC3*) from the red, and six (*CD2, CD247, TGFBR3, ARL4C, KLRK1, MATK*) from the black module (**Figure 5, Table S8**). For qPCR validation, available samples from HIV-negative TBM patients (n=132) were used to evaluate 11 hub genes selected from the blue and brown modules. 8/11 hub genes were found to be significantly associated with mortality, as determined by univariate Cox regression (**Table 3**). Three genes (FCAR, PGD, ETS2) were also found to be associated with mortality, but in the reverse direction.

**Figure 5.**
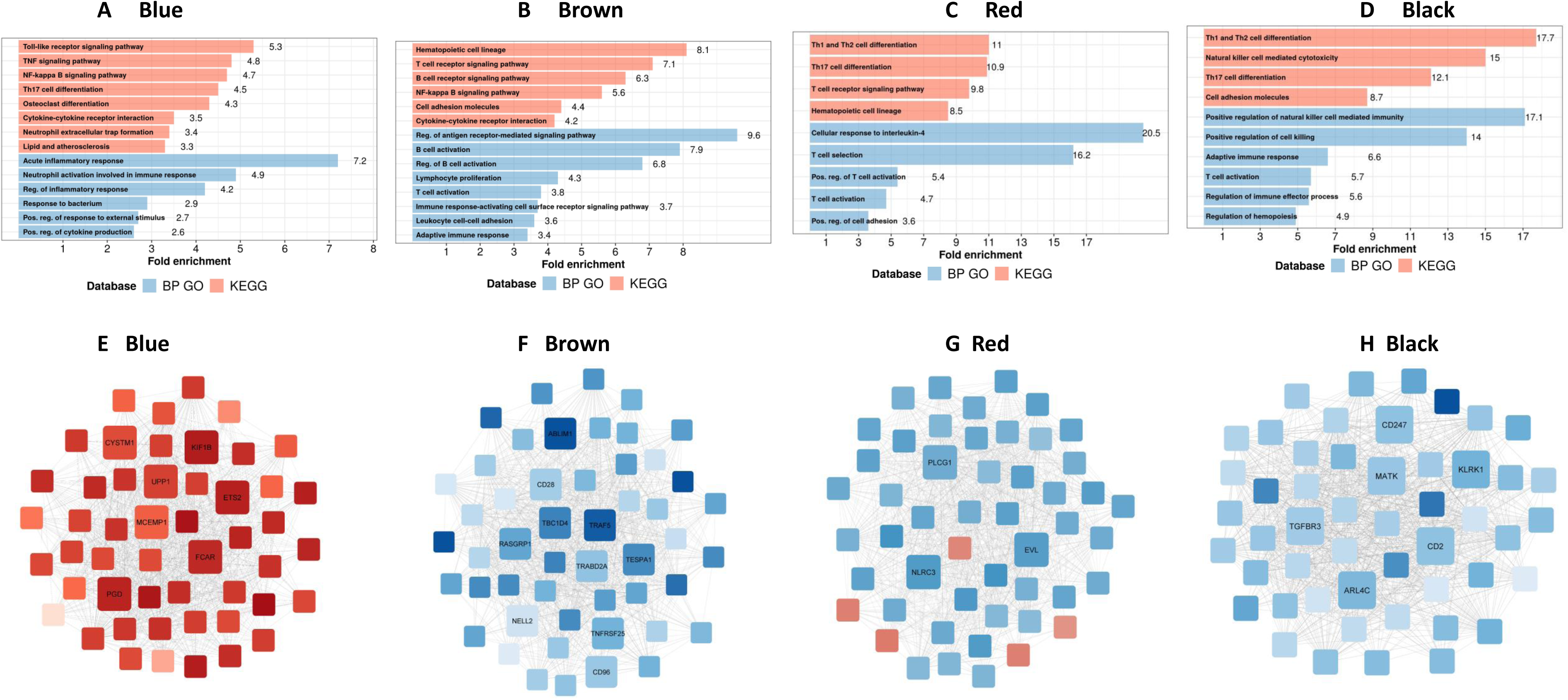
Biological processes, pathways and hub genes of validated modules associated with mortality. **(A-D)** showed biological processes and pathways identified in four mortality associated modules: blue, brown, red and black module, by over representation analysis (ORA). Bar plots show the top representative GO biological processes or KEGG pathways. The bars indicates biological processes or pathways having ORA FDR < 0.05 and size corresponding to fold enrichment calculated as the ratio of gene number of pathway in the input list divided by the ratio of gene number of the pathway in reference. **(E-H)** showed gene co-expression networks and hub genes of blue, brown, red and black module, respectively. Each node represents one gene. Each edge represents the link between two genes. Hub genes were shown by bigger nodes and bold text. The gradient color of node corresponds to its HR to mortality, with red indicating HR > 1, and blue HR < 1.

**Table 3.** Validation of hub genes in PCR validation cohort.

| No. | Genes | RNA-seq cohort<br>HIV-negative TBM (n=207) |  |  | qPCR validation cohort<br>HIV-negative TBM (n=132) |  |  |
| --- | --- | --- | --- | --- | --- | --- | --- |
|  |  | HR | 95%CI | p value | HR | 95%CI | p value |
| 1 | MCEMP1 | 1.93 | 2.37, 1.57 | 4.07E-10 | 1.74 | 2.37, 1.27 | 4.99E-04 |
| 2 | FCAR | 2.37 | 3.30, 1.70 | 3.52E-07 | 0.59 | 0.87, 0.40 | 7.87E-03 |
| 3 | ETS2 | 2.65 | 3.81, 1.85 | 1.24E-07 | 0.27 | 0.44, 0.16 | 3.27E-07 |
| 4 | PGD | 2.65 | 3.81, 1.84 | 1.47E-07 | 0.18 | 0.34, 0.09 | 2.69E-07 |
| 5 | NELL2 | 0.56 | 0.71, 0.45 | 6.08E-07 | 0.80 | 0.96, 0.66 | 1.53E-02 |
| 6 | TRABD2A | 0.43 | 0.58, 0.31 | 4.94E-08 | 0.64 | 0.8, 0.51 | 1.05E-04 |
| 7 | TRAF5 | 0.31 | 0.49, 0.2 | 4.52E-07 | 0.47 | 0.61, 0.36 | 1.66E-08 |
| 8 | CD28 | 0.57 | 0.76, 0.42 | 1.42E-04 | 0.60 | 0.76, 0.47 | 2.83E-05 |
| 9 | TESPA1 | 0.36 | 0.52, 0.25 | 2.74E-08 | 0.65 | 0.89, 0.48 | 7.35E-03 |
| 10 | ABLIM1 | 0.31 | 0.49, 0.2 | 2.21E-07 | 0.47 | 0.63, 0.36 | 1.55E-07 |
| 11 | RASGRP1 | 0.45 | 0.67, 0.31 | 6.10E-05 | 0.59 | 0.77, 0.45 | 9.77E-05 |

Association of the hub genes with three-month mortality using a univariate Cox regression model. Hazard ratio (HR), 95% CI of HR and p-value from the Cox regression model were presented in the table. HR or 95% mean per increase 1 unit of log2 normalized expression of gene in RNA-seq cohort or decrease 1 unit of cycle threshold in qPCR validation cohort.

We next examined patterns of shared and distinct gene expression of some hub across the four cohorts (healthy controls, PTB, and TBM with and without HIV-infection) to reveal disease progression and pathogenesis of different TB forms (**Figure 6**). There were two upregulated genes from the acute inflammation module (*FCAR* and *MCEMP1*) and six downregulated genes (*NELL2, TRABD2A, PLCG1, NLRC3, CD247* and *MATK*) from the other three adaptive immunity modules. The patterns of up and downregulation, relative to healthy controls, were similar in PTB and TBM, although tended to be more pronounced in those with TBM as well as those with HIV (**Figure 6**).

**Figure 6.**
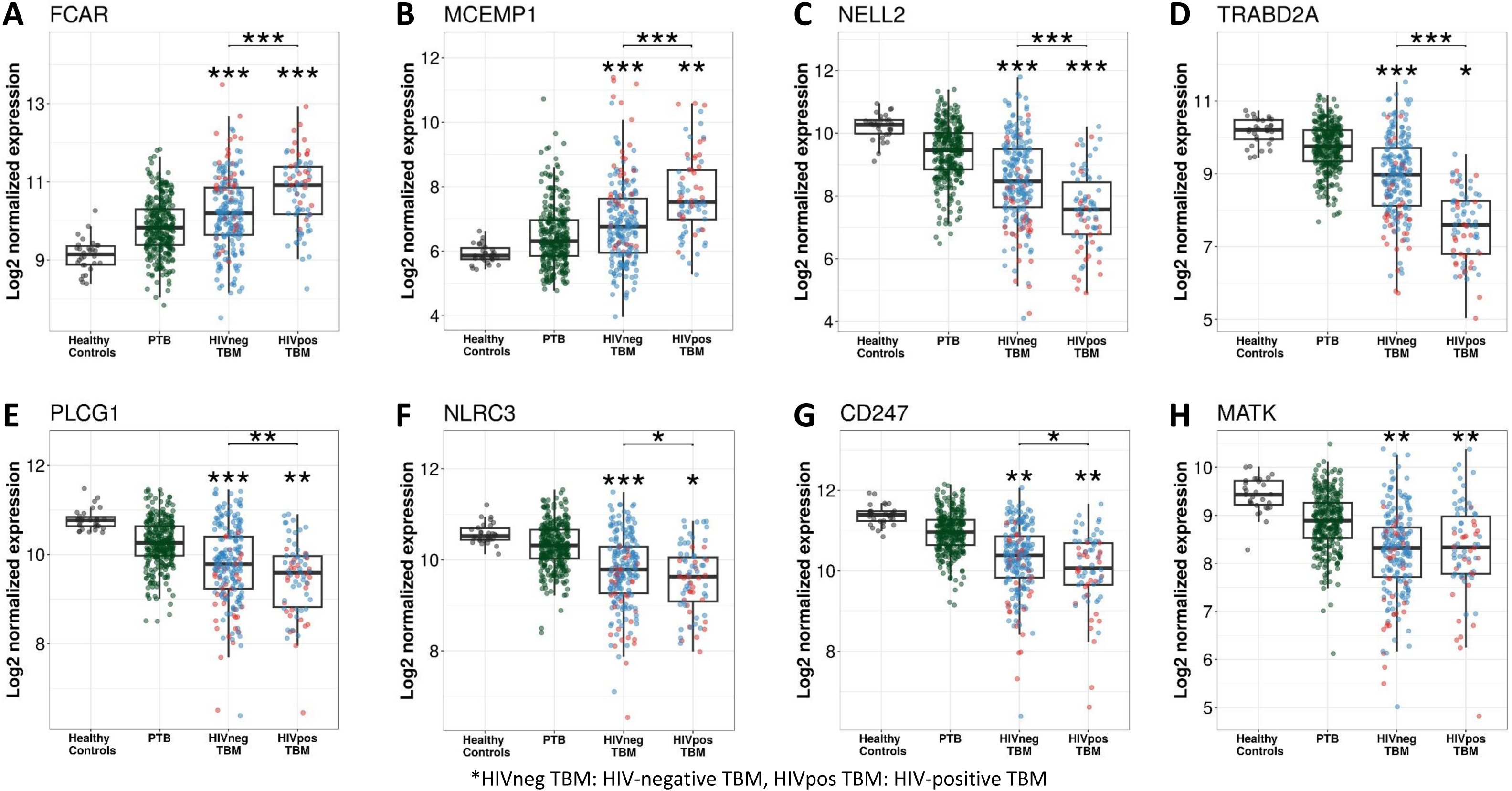
Gene expression of representative hub genes in healthy controls (n=30), PTB (n=295), HIV-negative TBM (n=207) and HIV-positive TBM (n=74) Each dot represents gene expression from one participant. **(A, B)** expression of FCAR and MCEMP1 hub genes from the blue module. **(C, D)** expression of NELL2 and TRABD2A hub genes from the brown modules. **(E, F)** expression of PLCG1 and NLRC3 hub genes from the red module. **(G, H)** expression of CD247 and MATK hub genes from the black module. The box presents median, 25^th^ to 75^th^ percentile and the whiskers present the minimum to the maximum points in the data. Comparisons were made between dead (red) with survival (blue) or between HIV-negative and HIV-positive TBM by Wilcoxon rank sum test with p-values displayed as significance level above the boxes and the horizontal bars (* < .05, ** < .01, *** <.001).

### Transcriptional immune pathways associated with TBM mortality

To better understand the biological functions of the five modules associated with TBM mortality, a gene set associated with mortality in each module was functionally annotated using known pathway databases, such as gene ontology (GO) and Kyoto Encyclopedia of Genes and Genomes (KEGG) ^19^. Pathway enrichment analysis was performed using fold enrichment to determine if the prevalence of genes in a pathway were different from that expected by chance. We also used ssGSEA enrichment scores based on gene expression ranking of genes in a particular pathway within a single individual, to show the activity of this pathway between death and survival TBM as well as across 4 cohorts. These analyses helped to identify the pathways linked to mortality within the gene modules.

In the blue module, we found TBM mortality was associated with upregulated inflammatory and innate immune response transcripts, particularly in pathways involved in the acute inflammatory response and the regulation of inflammatory responses, including responses to bacteria and neutrophil activation. KEGG pathway analysis suggested this signal was associated with transduction and immune system pathways, such as toll-like receptor signaling, TNF signaling, NF-kappa B signaling and neutrophil extracellular trap formation (**Figure 5, Table S9**).

We did not find any pathway significantly associated with mortality in the upregulated cyan gene module, although in the hierarchical clustering the cyan module was highly correlated with the blue - inflammatory response module (**Figure 4**). In contrast, analyses of the down-regulated brown, red and black modules highlighted pathways involved in adaptive immunity, predominantly those mediated by lymphocytes. These included downregulations of lymphocyte proliferation, T cell activation, B cell activation, natural killer cell mediated cytotoxicity and their signaling pathways such as antigen receptor-mediated, T and B cell receptor (**Figure 5**, **Table S9**).

We also investigated the association between pathways known to be important to TB pathogenesis and mortality (**Figure 7**). Looking at those who died vs. survived amongst HIV-positive TBM, there was little difference between the groups with respect to inflammatory response, cytokine signaling, and icosanoid and inflammasome activation. This was also the case for HIV-negative individuals. However, interferon and TNF pathways were more active in those with HIV, and TNF signaling expression was significantly higher in HIV-negative adults who died rather than survived.

**Figure 7:**
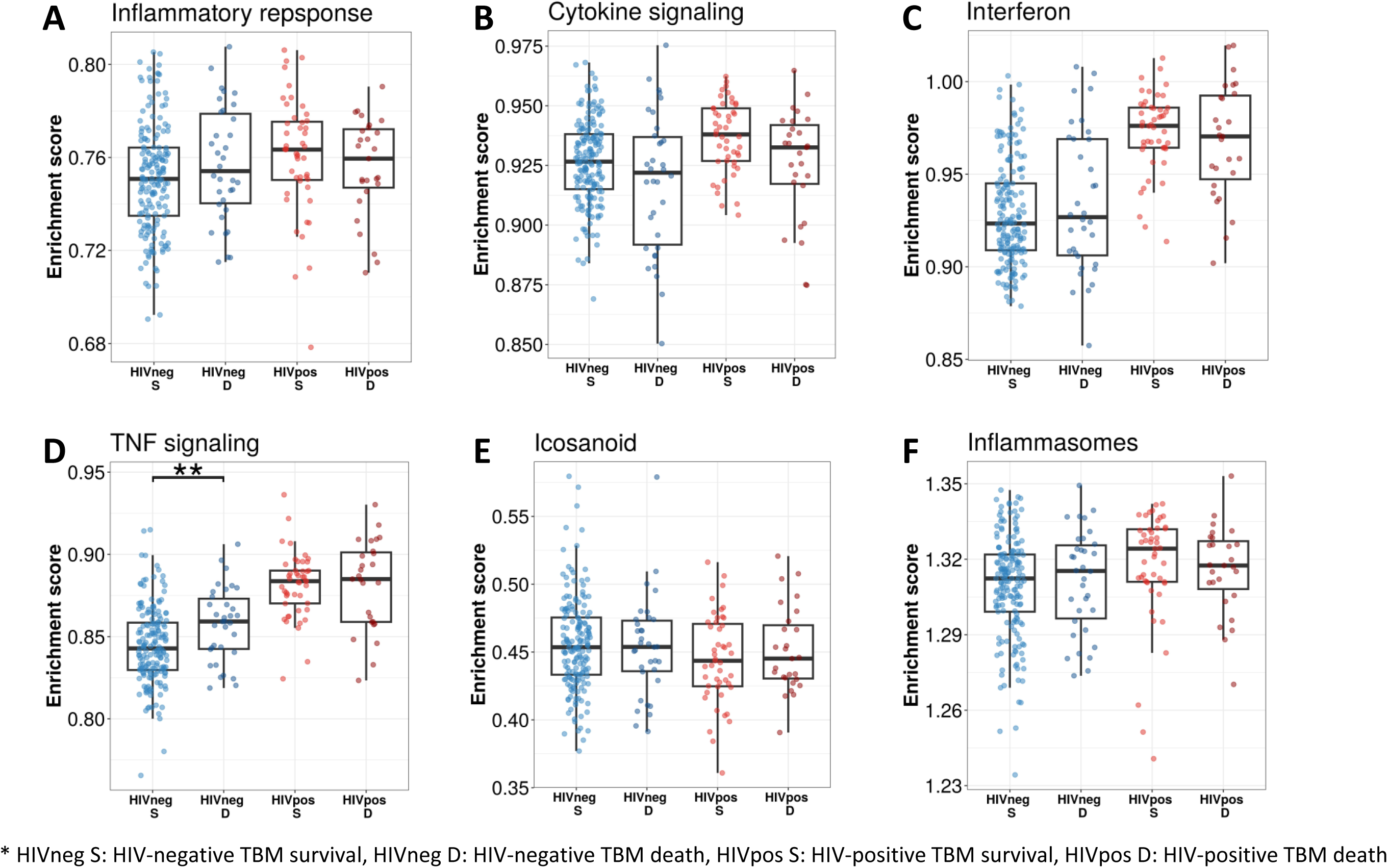
Relationship between known pathways associated with TBM pathogenesis and mortality. Enrichment scores of known immune pathways associated with TBM pathogenesis. Gene list of these pathway were depicted in additional file 1, table S5. Pathway enrichment scores were calculated using single sample GSEA algorithm (ssGSEA) (Barbie DA, et al 2009). Each dot represents one participant. The box presents median, 25^th^ to 75^th^ percentile and the whiskers present the minimum to the maximum points in the data. The comparisons were made between survival and death using Wilcoxon rank sum test. Only significant results are presented with * < .05, ** < .01, *** <.001.

### HIV influence on modules, hub genes, and pathways associated with TBM mortality

Transcriptional profiles associated with TBM mortality stratified by HIV status are shown in **Figure 3**. In HIV-negative individuals, 786 (3.9%) genes were differentially expressed, whereas in HIV-positive the number was 1620 (8.1%) genes (**Figure 3, Table S3 & S4)**. We hypothesized that host transcriptional signatures associated with TBM mortality differ according to HIV status. To test this hypothesis, we constructed gene co-expression networks from all genes in HIV-negative (n=207) and HIV-positive (n=74) individuals separately, then performed consensus gene co-expression network analysis (**Figure S4**). Modules were identified with colors and to discriminate them from the modules linked to mortality alone we added an annotated function on the module name. We focused on modules that failed to form consensus associations with mortality due to opposite associations in the two cohorts.

Sixteen gene co-expression modules (**Figure 8**, **Table S10**), ranging from 60 to 958 genes, were obtained from the HIV-negative and positive cohorts. Of these, 12 modules formed consensus association with mortality (**Figure 8**). Of the 4 modules which failed to form consensus association, two modules were significantly associated with mortality (FDR<0.05). The blue-angiogenesis module was up-regulated in death in HIV-positive adults (HR: 1.7) and the yellow-extracellular matrix organization (EMO) module was down-regulated in death in HIV-negative adults (HR: 0.31) (**Figure 8**).

**Figure 8.**
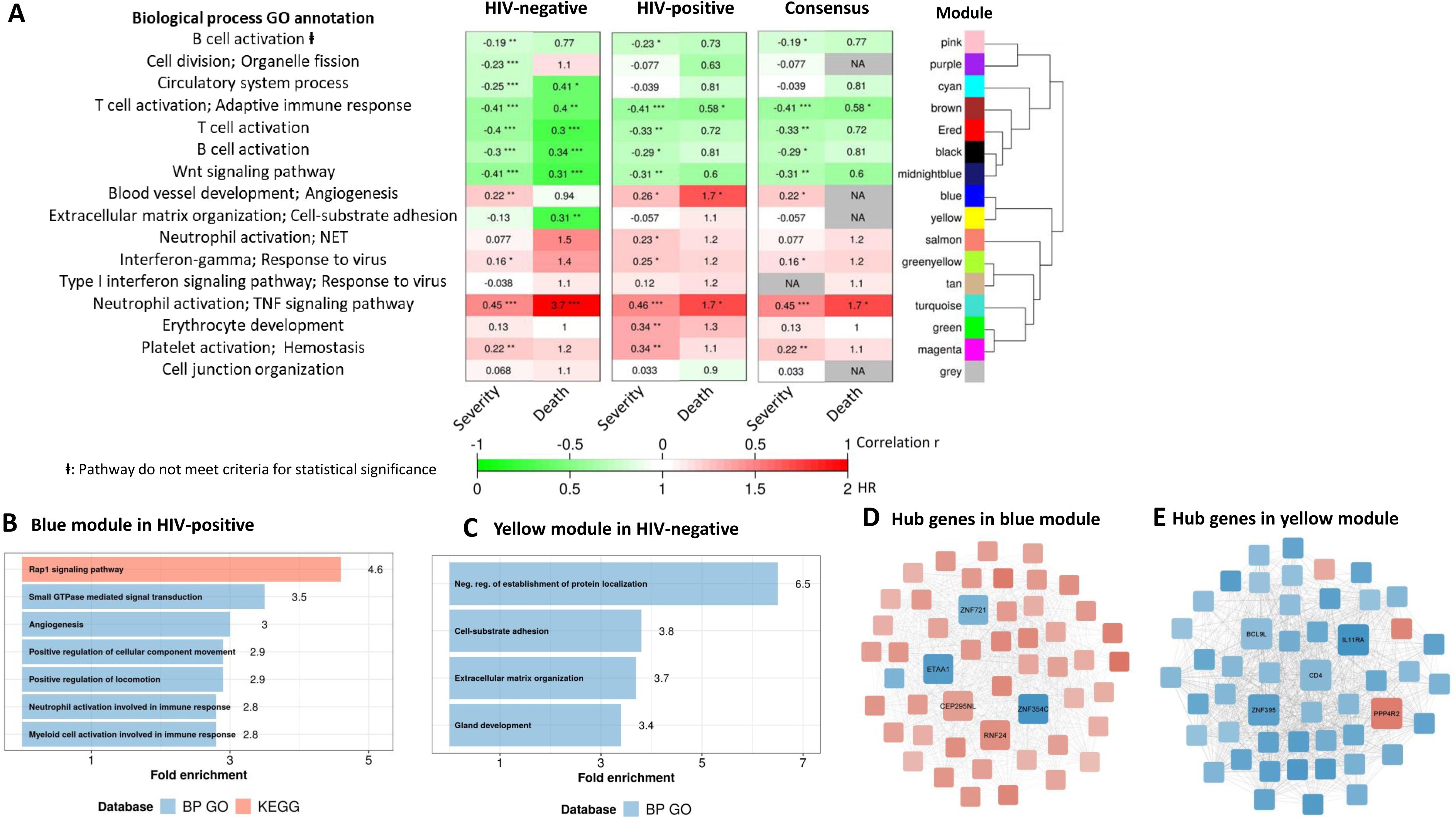
Consensus transcriptional modules associated with TBM mortality stratified by HIV-infection. **(A)** Associations between 16 consensus WGCNA modules with two clinical phenotypes TBM severity (MRC grade) and mortality in HIV-negative (n=207) and HIV-positive (n=74) TBM participants, and their associated BP Gene ontology or KEEG database. The heatmap showed the association between modules and the phenotypes, with Spearman correlation r for MRC grade and hazard ratio per increase 1 unit of PC1 of module (HR) for mortality in HIV-positive and HIV-negative cohorts. The consensus sub-panel presented associations of the consensus modules and clinical phenotypes with same trend detected in both HIV cohorts, otherwise were annotated with missing (NA) values. False discovery rate (FDR) corrected using Benjamini & Yekutieli procedure, with significant level denoted as * < .05, ** < .01 and *** <.001. Gradient colors were used to fill the cell with green indicating negative r or HR < 1, red color indicating positive r or HR > 1. The order of modules was based on hierarchical clustering using Pearson correlation distance for module eigengene. It is noted that these consensus modules were not identical to the identified modules in the primary analysis in Figure 1. **(B-C)** Functional enrichment analysis of HIV-positive pathway (blue module) and HIV-negative pathway (yellow module), respectively. **(D-E)** Gene co-expression network of blue and yellow modules. Each node represents one gene. Each edge represents the link between two genes. Hub genes were shown by bigger nodes with bold text. The gradient color of node corresponds to its HR to mortality, with red indicating HR>1, and blue HR<1.

Hub genes associated with mortality, which are highly correlated with other genes in the module, were identified in the two modules. Five hub genes, with three downregulated (*ZNF354C, ZNF721* and *ETAA1*) and two upregulated (*CEP295NL* and *RNF24*), were identified in the blue-angiogenesis module. Other five hub genes, with four downregulated (*IL11RA*, *CD4*, *ZNF395* and *BCL9L*) and one upregulated (*PPP4R2*), were identified in the yellow-EMO module (**Figure 8, Table S11**). Expression of some hub genes across the four cohorts showed the patterns of up (*RNF24*) and downregulation (*ZNF721, BCL9L* and *IL11RA*), and relative to healthy controls they were similar in PTB and TBM. Expression of *RNF24* and *ZNF721* genes were significantly associated with death in HIV-positive adults, whereas expression of *BCL9L* and *IL11RA* genes were significantly associated with death in HIV-negative adults (**Figure S5**).

Pathway enrichment analysis showed genes in the blue-angiogenesis module were significantly enriched for angiogenesis or blood vessel development, leukocyte and neutrophil activation, and signal transduction pathways (**Figure 8**, **Table S12**). Gene expression in the yellow-module was strongly enriched in extracellular matrix organization, cell-substrate adhesion and protein localization pathways (**Figure 8**, **Table S12**). Hierarchical clustering of modules indicated that these two modules were highly correlated (**Figure 8**) suggesting they share similar functions, which appeared in some pathways such as angiogenesis and extracellular matrix organization.

In addition, almost all of the pathways significantly associated with TBM mortality were similar in HIV-negative and positive cohorts, confirming that these important pathways were common to all TBM (**Figure S6**). However, pathway enrichment scores were higher in HIV-positive than in HIV-negative, especially for the TNF signaling pathway and TNF transcripts (**Figure 9**). Death in HIV-negative TBM was associated with increased enrichment of these pathways, but with less TNF transcript compared to HIV-positive individuals. The enrichment scores indicated mortality was strongly associated with downregulation of adaptive immune responses, including T cell activation, T and B cell receptor signaling pathways, with little impact of HIV status on these pathways (**Figure 9**). These data suggest an inadequate adaptive immune response contributes to disease pathogenesis and mortality in all those with TBM, regardless of HIV status.

**Figure 9.**
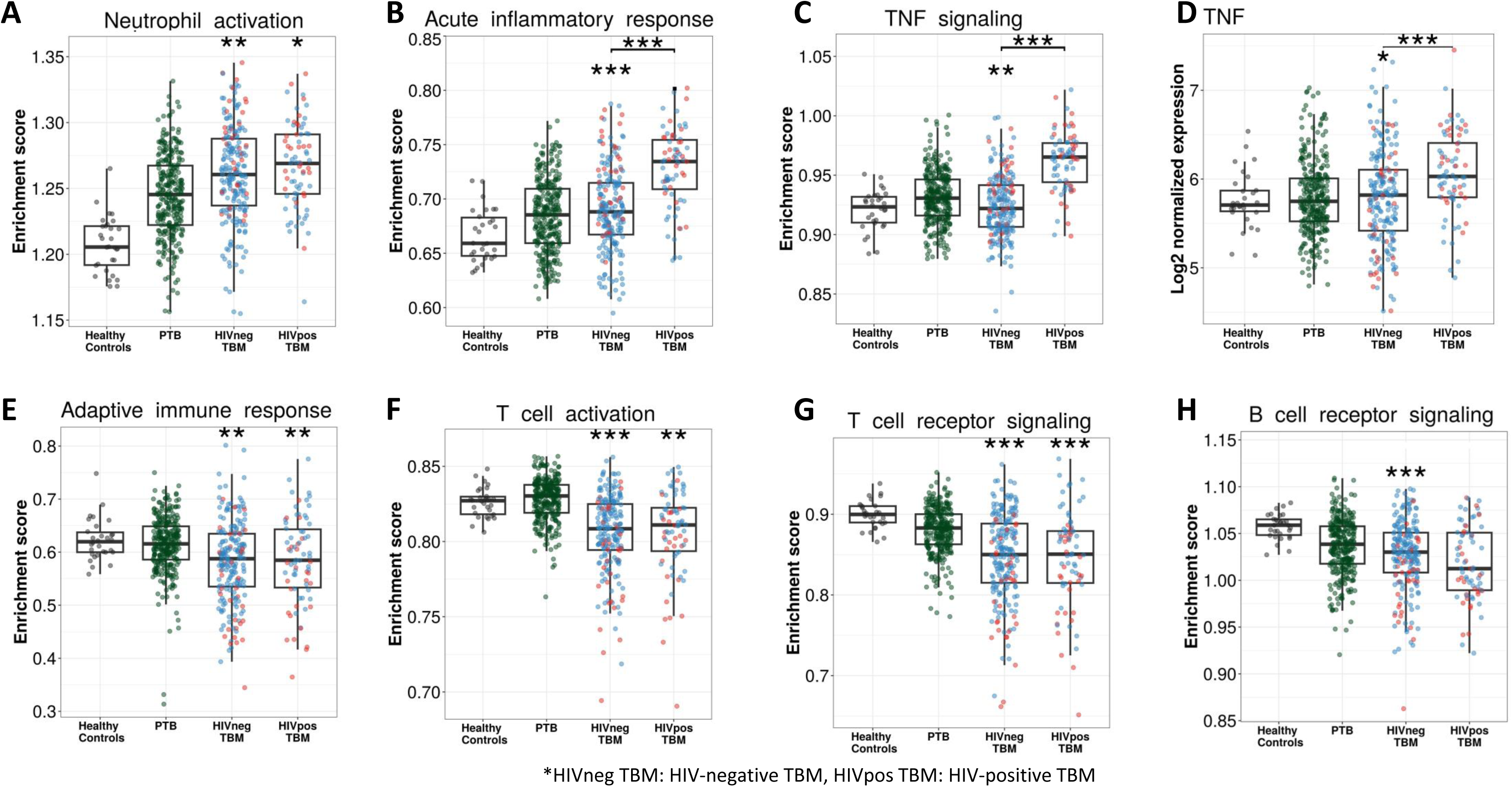
Enrichment score of immunity pathways in healthy controls (n=30), PTB (n=295), HIV-negative TBM (n=207) and HIV-positive TBM (n=74) Pathway enrichment scores were calculated using single sample GSEA algorithm (ssGSEA) (Barbie DA, et al 2009). Each dot represents one participant. **(A-C)** showed box-plots depicting enrichment scores of the innate immunity pathways from the blue module. **(E-H)** enrichment scores of the adaptive immunity pathways from the red and brown modules. **(D)** normalized expression of TNF. Gene list of these pathway were depicted in additional file 1, table S5. The box presents median, 25^th^ to 75^th^ percentile and the whiskers present the minimum to the maximum points in the data. Comparisons were made between dead (red) with survival (blue) or between HIV-negative and HIV-positive TBM by Wilcoxon rank sum test with p-values displayed as significance level above the boxes and the horizontal bars, respectively (* < .05, ** < .01, *** <.001).

### Predictors of TBM mortality

We aimed to identify baseline gene signatures that might help predict death or survival from TBM. We selected potential gene predictors from 26 common and 10 HIV-specific hub genes; these hub genes represented the dominant modules associated with mortality. Three predefined clinical factors known to be associated with outcome (age, MRC grade and CSF lymphocytes) were also used for candidate predictor selection. Using a multivariate Cox elastic-net regression model for 39 predictors, six predictors were selected for HIV-negative TBM (MRC grade, age, CSF lymphocyte count, gene set 1: *MCEMP1*, *TRABD2A* and *CD4*), and six predictors were selected for HIV-positive TBM (MRC grade, age, CSF lymphocyte count, gene set 2: *NELL2*, *MCEMP1* and *ZNF354C*) (**Table S13**).

Gene expression and association with mortality of five distinct genes in gene sets 1 and 2 were presented in **Figure S7**. By combining the two gene sets above and reducing genes in the same module, we generated gene set 3 (*MCEMP1*, *TRABD2A*, *CD4* and *ZNF354C*) and gene set 4 (*MCEMP1*, *NELL2*, *ZNF354C* and *CD4*). These two gene sets, with and without clinical factors (age, MRC grade and CSF lymphocytes), were tested for their predictive performance in HIV-negative and HIV-positive RNA-seq cohorts together with a reference model ^15^. Model 7 using gene set 4 and clinical factors outperformed other gene sets and reference model, with the best overall model performance (lowest optimism-corrected Brier score, 0.11 and 0.14 for HIV-negative and HIV-positive, respectively) and the best discriminant performance (highest optimism-corrected AUC 0.80 and 0.86 for HIV-negative and positive) (**Table 4**, **Figure S8**).

**Table 4.** Comparison of gene signatures in distinguishing survival and death in TBM prognostic models.

| **A. RNA-seq cohorts** |  |  |  |  |  |  |  |
| --- | --- | --- | --- | --- | --- | --- | --- |
| **No.** | **Predictor set** | **All TBM**<br>**n=281** |  | **HIV-negative TBM**<br>**n=207** |  | **HIV-positive TBM**<br>**n=74** |  |
|  |  | **AUC** | **Brier score** | **AUC** | **Brier score** | **AUC** | **Brier score** |
| 1 | **Reference model** (PMID: 29029055) | 0.78 | 0.14 | 0.77 | 0.12 | 0.82 | 0.18 |
| 2 | **Gene set 1** (MCEMP1, TRABD2A, and CD4) | 0.78 | 0.14 | 0.78 | 0.11 | 0.65 | 0.23 |
| 3 | **Gene set 2** (MCEMP1, NELL2 and ZNF354C) | 0.78 | 0.14 | 0.77 | 0.11 | 0.75 | 0.20 |
| 4 | **Gene set 3** (MCEMP1, TRABD2A, CD4 and ZNF354C) | 0.78 | 0.14 | 0.77 | 0.11 | 0.73 | 0.21 |
| 5 | **Gene set 4** (MCEMP1, NELL2, CD4 and ZNF354C) | 0.79 | 0.14 | 0.77 | 0.11 | 0.75 | 0.20 |
| 6 | **Gene set 3** and clinical risk factors | 0.82 | 0.13 | 0.80 | 0.11 | 0.84 | 0.15 |
| 7 | **Gene set 4** and clinical risk factors | 0.82 | 0.13 | 0.80 | 0.11 | 0.86 | 0.14 |
| **B. qPCR validation cohort** |  |  |  |  |  |  |  |
| **No.** | **Predictor set** | **HIV-negative TBM (n=132)** |  |  |  |  |  |
|  |  | **AUC** |  |  | **Brier score** |  |  |
| 8* | MCEMP1, NELL2, ZNF354C and clinical predictors | 0.92 |  |  | 0.12 |  |  |

Clinical risk factors were age, MRC grade and CSF lymphocytes<sup>14</sup>. The prediction models for three-month mortality were based on multivariable logistic regression models with top-hit genes and clinical risk factors. Area under the curve (AUC) and Brier score were corrected for optimism using internal bootstrap resampling over 1,000 iterations to evaluate the model performance. The Brier score is an overall performance measure, calculated as the mean squared difference between the predicted probability and the actual outcome, with smaller values indicating superior model performance.
\* CD4 data is unavailable for the analysis.

Given that gene expression could reflect cellular composition changes of the peripheral blood, with neutrophils being the most abundant sub-population, we performed a sensitivity analysis including blood neutrophil count as a potential gene predictor in a multivariate Cox elastic-net regression model. In the sensitivity analysis, all six predictors for either HIV-negative TBM or HIV-positive TBM were repeatedly selected (**Table S14**). We evaluated the predictive values of blood neutrophil alone and in combination with identified gene sets, but adding neutrophils did not improve the overall performance of the predictive models.

Next, we evaluated the prognostic signature of model 7 in another sample set of 132 HIV-negative TBM patients with gene expressions measured by qPCR. Only three genes (*MCEMP1, NELL2* and *ZNF354C*) from gene set 4 were used because qPCR data of *CD4* was not available. The analysis demonstrated good predictive performance in model 8, with an optimism-corrected AUC 0.92 and optimism-corrected Brier score 0.12 (**Table 4**, **Figure S8**). Despite using three-gene signature in the validation, the results still confirmed the ability to predict TBM mortality in HIV-negative patients.

## Discussion

The biological pathways involved in pathogenesis of TBM are unclear. In general, previous studies investigating TBM pathogenesis have been small, testing for relatively small numbers of selected genes or molecules, and have been unable to take an unbiased and broader view of the inflammatory response. This study investigated the pathways associated with death from TBM at a whole-genome transcriptome level in whole blood, characterizing a global dysregulation in immune responses, including inflammation, and revealing specific functional pathways and hub genes involved in TBM and the mechanisms leading to death.

We sought to understand better the pathogenesis of TBM by identifying pretreatment blood transcriptional gene modules associated TBM disease severity and three-month mortality. Four out of five identified modules were involved in immunological functions, indicating multiple functional pathways of systemic immunity are involved in the pathogenesis of TBM. In particular, mortality was strongly associated with increased acute inflammation and neutrophil activation, and decreased adaptive immunity and T and B cell activation. Whilst there appeared to be many common pathways involved in TBM mortality in HIV-positive and negative individuals, there were differences: death was associated with increased expression of angiogenesis genes in HIV-positive adults, and with TNF signaling and down regulated extracellular matrix organization in HIV-negative adults. We also identified a four-gene signature as a potential host response biomarker for mortality, regardless of HIV status.

TBM mortality was associated with increased acute inflammatory responses, the regulation of inflammatory responses, and neutrophil activation. Previous blood transcriptome studies have shown that IFN-inducible neutrophil-driven transcripts were over-expressed in blood neutrophils from active TB patients compared to those with latent TB ^20,21^. Another study described that TBM adults who developed IRIS during treatment, also had significantly more abundant neutrophil-associated transcripts that preceded the development of TBM-IRIS ^5^. Our own earlier studies have also suggested a role for neutrophils in TBM pathogenesis, with high pretreatment CSF bacterial loads being correlated with high neutrophil numbers in both CSF and blood and more frequent new neurological events or paradoxical inflammatory complications ^8^.

Taken together with previous studies, our findings support an important role for over-activation of neutrophil-mediated inflammatory responses in TBM pathogenesis and its lethal complications. Looking further into specific pathways, TBM mortality was neither associated with inflammatory response and cytokine signaling pathways in general, nor with interferon or inflammasome activation. However, increased transcripts in some specific immunity pathways, including TNF signaling, Toll-like receptor, NF-kappa B and neutrophil extracellular trap formation, were associated with TBM mortality. These pathways are involved as activators or effectors in the process of neutrophil activation and regulation, leading to an exacerbated inflammatory response ^22–24^.

We found blood transcriptional responses of T cells and B cells were under-expressed in those who died from TBM, indicating an impairment in adaptive immunity in fatal disease. A reduction of activities in both T cell and B cell receptor signaling pathways in death were identified, independent of over-expression of neutrophil-mediated immune responses, indicating that multiple functional pathways influence TBM mortality. TBM pathogenesis is known to be associated with T cell impairment. Previous studies have shown lower numbers of T cells, reduced ability to respond to *Mycobacterium tuberculosis* antigens or reduced expression of activation markers and cytokine production in TBM compared to PTB and healthy individuals ^10,25,26^. This impaired T cell function has correlated with disease severity and poor clinical outcomes in participants with PTB and TBM ^10,27,28^.

B cells and antibodies also influence humoral immunity against *Mycobacterium tuberculosis*. Studies have shown a decreased memory B cell proportion, and lower levels of IgG, IgM antibodies and Fcγ receptors binding capacity in plasma in those with active lung TB compared to those with latent TB or healthy volunteers ^29,30^. But little is known about the role of B cells in TBM. The observed association between TBM mortality and decreased transcriptional responses in B cell activation and B cell receptor signaling pathways suggest an unanticipated role for B cells and humoral immunity in TBM pathogenesis that needs further investigation.

Transcriptomic profiles from four cohorts, including healthy controls and PTB, provide a broad view of host responses in different TB clinical forms. Our data showed common transcriptional pathways and genes between PTB and TBM. A range of immune responses, involving in inflammation, cytokines, interferon, inflammasome and neutrophil signaling pathways, were activated in both PTB and TBM, but with significantly greater activation in HIV-positive TBM than HIV-negative TBM. This finding aligns with our previous data showing a dyregulated hyper-inflammation in HIV-associated TBM, with significantly higher CSF cytokine concentrations than in those without HIV infection ^7^. These data suggest that different forms of TB are associated with similar inflammatory responses, but with different degrees of host responses, exemplified by hyper-inflammation in those with HIV.

The HIV-driven differences in TBM-associated inflammation appear sufficient to influence response to adjunctive anti-inflammatory therapy with corticosteroids. An earlier randomized controlled trial of corticosteroids in 545 predominantly HIV-negative Vietnamese adults showed they significantly improved survival ^31^. Our group recently completed a similar sized trial exclusively in HIV-positive adults without any clear benefit upon mortality ^32^. Therefore, the differences in gene expression associated with TBM mortality in the current study – with angiogenesis activation linked to HIV-positive TBM mortality only, for example – may provide an explanation for the poor response and offer alternative therapeutic strategies. Recent studies have shown that angiogenesis is induced by *Mycobacterium tuberculosis* infection, which then contributes to inflammation, tissue damage and is correlated with disease severity ^33^.

Developing prognostic models for TBM is important for guiding clinical decision making and improving outcomes. Several studies have developed and validated prognostic models for TBM using clinical, laboratory and radiological variables. These models have demonstrated moderate to high accuracy in predicting mortality and functional outcomes from TBM patients ^15,34–36^. In this study, we used blood transcriptional signatures and co-expression network analysis to identify module-representative hub genes. We identified a four-gene set at the start of treatment (*MCEMP1*, *NELL2*, *ZNF354C* and *CD4*) whose expression strongly predicted three-month TBM mortality. Our prognostic models combining this four-gene host response in blood and three routine clinical predictors achieved very good performance, with AUC 0.80 and 0.86 for HIV-negative and HIV-positive individuals with TBM, respectively. This is proof-of-concept that whole blood RNA host response might be a useful pre-treatment biomarker to predict early TBM mortality. Although further investigation and validation is needed, we have identified potential gene candidates for future development as prognostic biomarkers.

In summary, we present a comprehensive and unbiased analysis of the gene transcripts in blood associated with TBM severity and mortality. Our data open a new window on TBM pathogenesis, with dysregulation in both innate and adaptive immune responses strongly associated with death from TBM. Furthermore, we have identified similarities and differences in the inflammatory responses associated with TBM in HIV-positive and negative adults, which may explain the different therapeutic effects of adjunctive corticosteroid treatment. We also revealed a 4-gene host response signature in blood that might represent a novel biomarker for defining those at highest risk of death, regardless of their HIV status. The signature needs validation in future cohorts.

## Materials and Methods

### Participants

We collected data and whole peripheral blood samples for transcriptomic profiling from adults (≥18 years) with TBM enrolled into two randomized controlled trials conducted in Vietnam. The two trials investigated whether adjunctive dexamethasone improves outcome from TBM and the protocols for both trials have been published ^37,38^. The ACT-HIV trial (NCT03092817) completed enrolment and follow-up of 520 HIV-positive adults with TBM in April 2023, with the results accepted for publication ^32^. The LAST-ACT trial (NCT03100786) completed enrollment of 720 HIV-negative adults with TBM in March 2023, with the last participants due to complete follow-up in March 2024.

In TBM RNA-seq cohorts, peripheral blood samples were taken from 281 trial participants: the first 207 consecutively enrolled HIV-negative participants from the LAST-ACT trial, and 74 randomly selected HIV-positive participants from the ACT-HIV trial. In qPCR TBM cohorts, 132 HIV-negative participants were randomly selected from remaining participants of LAST-ACT trial with an enrichment for non-survival cases. The samples were taken at enrollment, when patients could not have received more than 6 consecutive days of two or more drugs active against *Mycobacterium tuberculosis*. All trial participants then received standard 4-drug anti-tuberculosis treatment for 2 months, followed by 3 drugs for 10 months, and were randomly allocated to dexamethasone or identical placebo for the first 6-8 weeks ^37,38^. The investigators remain blind to the treatment allocation until the last participant completes follow-up and the database has been locked. Therefore, an analysis of the direct influence of corticosteroids on inflammatory response and outcome is not included in the current study. To adjust for corticosteroid effect, treatment outcome and effect were blinded in analyses by extracting only the fold change difference between survival and death in the linear regression model, in which gene expression was outcome and survival and treatment were covariates.

Peripheral blood samples were taken for transcriptional profiling from two other cohorts: 295 adults with PTB and 30 healthy controls. The 295 PTB was enrolled in Pham Ngoc Thach hospital and District TB Units in Ho Chi Minh city, randomly selected from HIV-negative participants from a prospective observational study of the host and bacterial determinants of outcome from PTB (n=900) due to the very low prevalence of HIV in the study population. Participants had culture-confirmed PTB, either drug susceptible TB or a new diagnosis of multidrug-resistant TB. All participants in this cohort had <7 days of anti-TB drugs at enrolment and did not have clinical evidence of extra-pulmonary TB. Healthy controls were enrolled in Hospital for Tropical Diseases in Ho Chi Minh city, from a prospective study for epidemiological characteristics of human resistance to *Mycobacterium tuberculosis* infection. Participants in this cohort were adults without signs or symptoms of TB nor history of TB contact within the last two years.

Ethics approval was obtained from the institutional review board at the Hospital for Tropical Diseases, Pham Ngoc Thach Hospital and the ethics committee of the Ministry of Health in Vietnam, and the Oxford Tropical Research Ethics Committee, UK (OxTREC 52-16, 36-16, 24-17, 33-17 and 532-22). All participants provided their written informed consent to take part in the study, or from their relatives if they were incapacitated.

### Study design

The objectives and cohorts used in this study are presented in **Figure 1** and a workflow of data analysis is shown in **Figure S10**. Briefly, blood transcriptional profiling was generated from four cohorts of 207 TBM HIV-negative, 74 TBM HIV-positive, 295 PTB and 30 healthy controls. To define transcriptional signatures associated with TBM mortality, data from all 281 TBM participants were used. To ensure reproducibility, in our main analysis we randomly split our TBM data into two datasets, a discovery cohort (n=142) and a validation cohort (n=139). After identifying gene modules associated with mortality, related functional pathways and hub genes were determined with more details of the data analysis provided in **Figure S10**. In a broader view, gene enrichment and expression from significant pathways and hub genes were illustrated across all four cohorts. Next, outcome prediction models were developed for TBM. Finally, the association of hub genes and outcome prediction then was validated in qPCR HIV-negative TBM cohort.

### Sample processing and RNA-seq

Whole blood samples, collected from participants at enrollment, were stored in PAXgene collection tubes at −80^0^C. RNA extraction and RNA-seq were done in 2 batches. Batch 1 was done in 2020 including 207 HIV-negative TBM, 31 HIV-positive TBM and 295 PTB. Batch 2 was done in 2022 including 43 HIV-positive TBM and 30 Healthy control. RNA samples were isolated using the PAXgene Blood RNA kits (Qiagen, Valencia, CA, USA) following the manufacturer’s instructions, except for an additional washing step before RNA elution. DNA was digested on columns using the RNase-free DNase Set (Qiagen, Valencia, CA, USA). Quality control of the RNA extraction was performed using the Epoch spec for quantity and quality, and Tapestation Eukaryotic RNA Screentape for integrity. Samples with RNA integrity number below 4 were exclude for further steps. RNA-seq was performed by the Ramaciotti Centre for Genomics (Sydney, Australia). One microgram of total RNA was used as input for each sample, using the TruSeq Stranded Total RNA Ribo-zero Globin kit (Illumina). Libraries were generated on the Sciclone G3 NGS (Perkin Elmer, Utah, USA) and the cDNA was amplified using 11 PCR cycles. Libraries were pooled 75 samples per pool and sequenced using NovaSeq 6000 S4 reagents at 2×100bp to generate about 30 million reads per sample.

### RNA-seq data quality control and pre-processing

Quality control and alignment were performed using an in-house pipeline modified from previously published practices for RNA-seq analysis ^39,40^ in linux command line. Briefly, the quality of the sequencing fastq files was analyzed using FastQC (v0.11.5) and poor quality samples were excluded from further analysis. Sequence reads were adapter and quality trimmed using Trimmomatic (v0.36), followed by duplicated optical read removal using BBMap (v38.79) tool. STAR aligner (v2.5.2a) was used to align the reads to the human reference genome (GRCh38 build 99) downloaded from Ensembl, allowing for maximum 2 mismatches in each 25 bp segment and a maximum of 20 alignment hits per read ^41,42^. The alignment results were sorted and indexed for downstream analyses as BAM format files. The aligned reads were further utilized to generate gene expression counts using FeatureCounts (v2.0.0) against the human reference annotation (GRCh38 build 99) ^43^. Next, 60,067 genes in the expression matrix were first normalized by variance stabilizing transformation method built in DESeq2 package in R ^44^. Subsequently, the batch effect was removed for 20,000 most variable genes using combat function in the SVA package with correcting for 2 RNA-seq batches ^45^. The results of batch effect removal were visualized by principle component analysis. The first component was plotted against the second component. The variation explained by RNA-seq batches was removed after using combat (**Figure S9**). All later analysis and data visualization were done used batch corrected data.

### Validation of hub genes by Microfluidic multiplex RT-qPCR

Whole blood samples were collected, stored and RNA extraction was performed as described in the previous section. The expression of housekeeping genes (*GAPDH* and *TMBIM6*) and other hub genes were evaluated by microfluidic RT-qPCR using Biomark™ 48.48 Complete Bundle with Delta Gene™ Assays and BioMark™ HD system (Fluidigm Corporation, South San Francisco, CA, USA) following manufacturer’s instructions with some optimized modifications. Briefly, 2 µL of total RNA with concentration of 50 ng/µL was reverse transcribed to cDNA. The specific target amplification (STA) of cDNA was used for 14 cycles of preamplification, then the STA products were treated with Exonuclease I (New England Biolabs, Ipswich, MA, USA) before 20-fold dilution. SsoFast™ EvaGreen® Supermix with Low ROX (Bio-Rad Laboratories, Hercules, CA, USA) was used in RT-qPCR before applying to IFC Controllers MX and BioMark™ HD system (Fluidigm Corporation, South San Francisco, CA, USA). The cycle threshold (CT) values of target hub genes were normalized by subtracting for CT of *GAPDH* before analysis.

### Statistical analysis

The primary outcome examined in this study was TBM three-month mortality. In the descriptive analysis, we summarized and tested association of patient characteristics with three-month mortality using univariate Cox regression analysis. We presented the proportion for binary variables and median (1^st^ and 3^rd^ interquartile range) for continuous variables.

To explore transcriptional profiles associated with mortality in TBM, differential expression analysis was performed on the 20,000 normalized gene expression matrix to find differentially expressed genes (DEGs). We constructed the contradicted matrix with survival and death status adjusted for covariates including: age, HIV status, dexamethasone treatment (LAST-ACT trial investigators remained blind), and sequencing batch. Linear models were used to assess DEGs using limma R package. Empirical Bayes moderated-t p-values were computed for each genes and Benjamini-Hochberg were used for correcting multiple testing (FDR). We defined the DEGs with the parameters (fold change > 1.5 and FDR < 0.05). To visualize samples clustering in the 4 cohorts, an unsupervised principal component analysis was performed on 20,000 normalized genes, and the first principle component was plotted against the second principle component. The 95% confidence ellipse was drawn for each group using multivariate t-distribution ellipse. To compare enrichment activity of interested pathways between death vs survival, TBM vs PTB or healthy controls, enrichment scores of these pathways were calculated for single patient using single sample Gene Set Enrichment Analysis algorithm (ssGSEA). Pairwise Wilcoxon rank sum test was used for comparison between any two cohorts.

In the primary analysis, we initially conducted *weighted gene co-expression network analysis* (WGCNA) on the whole-blood transcriptomic profiling of the discovery cohort to identify clusters of genes (or ‘modules’) in the gene co-expression network among host transcriptomic genes. We then performed a *network modules preservation analysis* to assess the reproducibility of these modules and the network in the validation cohort. Subsequently, we conducted a *module-trait association analysis* to identify modules associated with the clinical traits of mortality and TBM severity. For the association analysis with mortality, we used a *Cox regression model* with the PC1 of the module as an independent variable. We adjusted the model for age, HIV status, and corticosteroid treatment (dexamethasone vs. placebo).

To analyze the association with TBM severity, we calculated the *Spearman correlation* between TBM severity (MRC grade) and the module’s PC1. We corrected for false discovery rate (FDR) for the dependent hypotheses by using the Benjamini-Yekutieli procedure. Target modules were defined as modules associated with mortality in both the discovery and validation cohorts at FDR < 0.05. Hub genes within each targeted module were identified based on 3 criteria: being protein coding gene, module membership cut off 0.85 (MM) and high rank of gene significance rank (GS). MM defined as correlation between individual gene and the module’s PC1. GS defined as −log_10_ p value association of gene with mortality using Cox regression model adjusted for age, HIV status and corticosteroid treatment. Genes were first filtered for protein coding function and MM above 0.85. Genes then was ranked based on its GS and top 20 hub genes were selected if number of genes in module above 500, otherwise top 10 genes were selected. Overlap hub genes between discovery and validation cohort were consider validated.

In the subsequent analysis, to determine the biological functions or processes potentially related to each module we conducted *overrepresentation analysis* (ORA) using ShinyGO v0.77 ^19^. Genes were first filters for association with three-month mortality at p < 0.05 and then were input into ShinyGO. We used both *GO and KEGG databases* for this analysis. We pre-specified significant pathways with overlap gene in pathway > 5 and FDR < 0.05.

Furthermore, to compare *enrichment activity* of resulting pathways between death vs survival, TBM vs PTB or healthy controls, enrichment scores of these pathways were calculated for single patient using ssGSEA algorithm ^46^. Pairwise Wilcoxon rank rum test were applied for comparison between 2 cohorts. To compare or visualize the difference of pathway effect on mortality between HIV-negative and HIV-positive TBM, *fold change enrichment* of interested pathways were calculated using Quantitative Set Analysis for Gene Expression (QuSAGE) method ^47^.

In the secondary analysis, we performed a *consensus analysis* to identify consensus patterns of co-expression networks between HIV-positive and HIV-negative conditions in our TBM cohort. We conducted similar association analyses for TBM severity and mortality as in the preceding WGCNA analysis for both the HIV-negative and HIV-positive cohorts. We visualized the results using a heatmap. We defined strongly associated modules in HIV-positive or HIV-negative cohort as HIV-positive-specific signals and HIV-negative-specific signals. Additionally, we functional enrichment ORA was performed for these consensus modules and the HIV-specific modules. Top 5 hub genes from HIV-specific modules were selected based on criteria described in the primary analysis, which included protein-coding genes and having high MM and high rank GS. GS in this analysis was calculated as −log_10_ p value association of genes with mortality using Cox regression model adjusted for age and corticosteroid treatment.

We then performed variable selection using a *multivariate elastic-net Cox regression model* to select the important predictors for HIV-negative and HIV-positive TBM prognosis separately. Candidate predictors in HIV-negative TBM consisted of the combination of common hub genes, HIV-negative specific hub genes and clinical biomarkers: age, TBM severity and CSF lymphocyte count. Similarly, for HIV-positive TBM, candidate predictors consisted of common hub genes, HIV-positive specific hub genes and clinical biomarkers. We performed the analysis with 1,000 bootstrap sampling with replacement. The chosen variables were 1) among the top 75% of selected variables and 2) one representative hub gene per module. Subsequently, we developed a prediction model based on a *logistic regression model* incorporating only the chosen variables for all TBM cohort or stratified by HIV-negative and HIV-positive cohort. We also assessed predictive performance of chosen variables in each cohort by 1000 times bootstrapping sampling approach and reported the overall model performance (*optimism-corrected Brier score*), the discrimination (*optimism-corrected AUC*), and calibration (*optimism-corrected calibration-slope*) of the developed model ^48^.

More details on performing ORA, SSGSEA, QuSAGE and WGCNA preservation and consensus analyses were provided in in the *Supplementary method document*.

## Acknowledgements

We would like to thank and the clinical staff who recruited patients into our study from Hospital for Tropical Diseases and Pham Ngoc Thach hospital, Ho Chi Minh city, Vietnam, and all participants.

## Supplementary Figures

**Figure S1.**
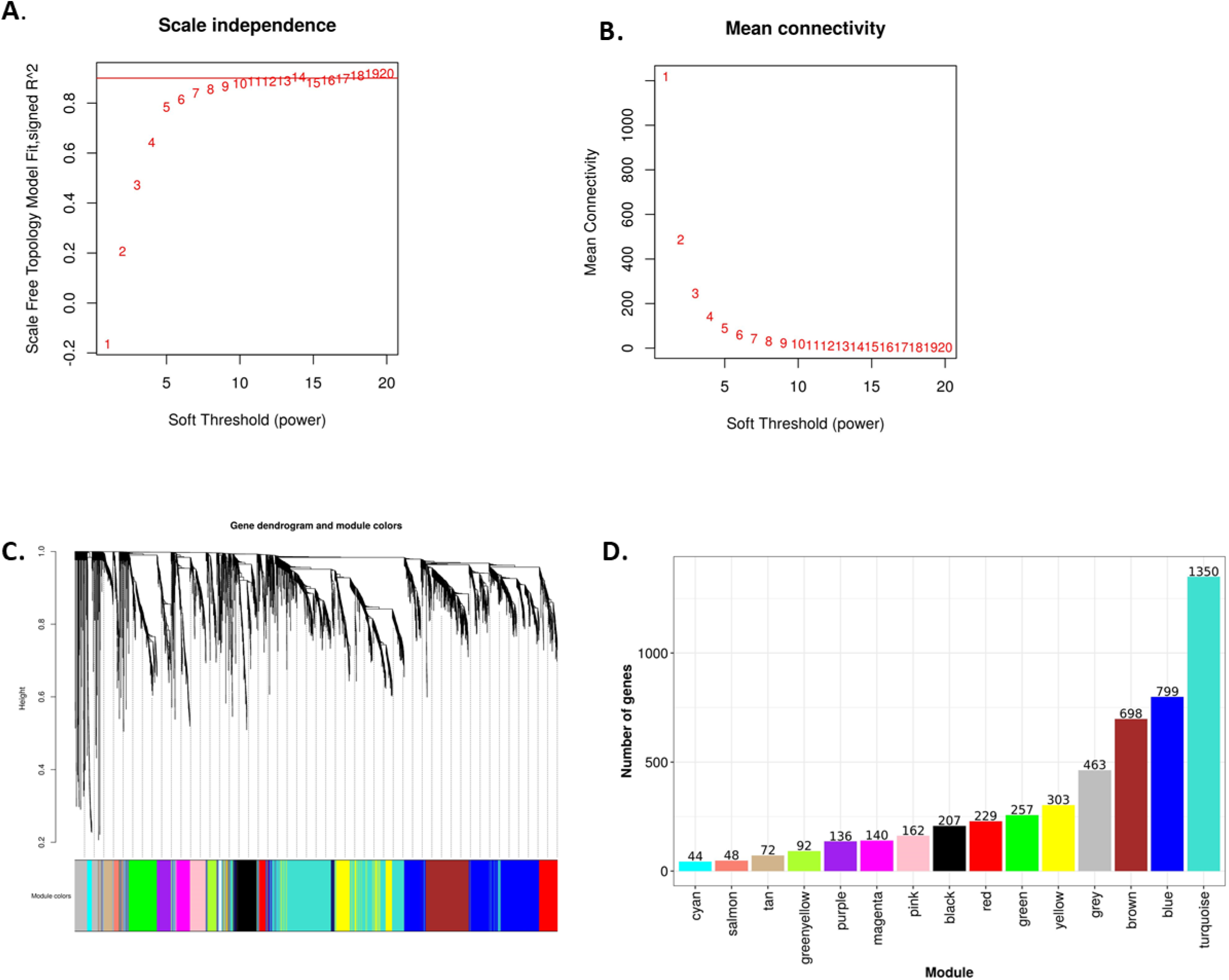
Construction of WGCNA in discovery cohort. Analysis of network topology for various soft-thresholding powers of the top 5,000 most variant genes based on the scale-free network model. **(A)** showed the scale-free fit index (y-axis) as a function of the soft-thresholding power (x-axis). The red horizontal line corresponds to *R*^2^ = 0.85 and soft-thresholding power *β*=8, which was chosen for the construction gene-expression network. **(B)** displayed the mean connectivity on the y-axis as a function of the soft-thresholding power on the x-axis. The adjacency matrix of the scale-free network between genes was determined as *A* = (*a_ij_*), where *a_ij_* = |*cor*(*gene_i_*, *gene_j_*|^*β*^. **(C)** displays the dendrogram resulted from the hierarchical clustering analysis using topological overlap of the adjacency matrix A served as a dissimilarity metric. Each cluster was referred as a module and assigned with a color. **(D)** showed bar-plots indicate the number of genes contained in each module.

**Figure S2.**
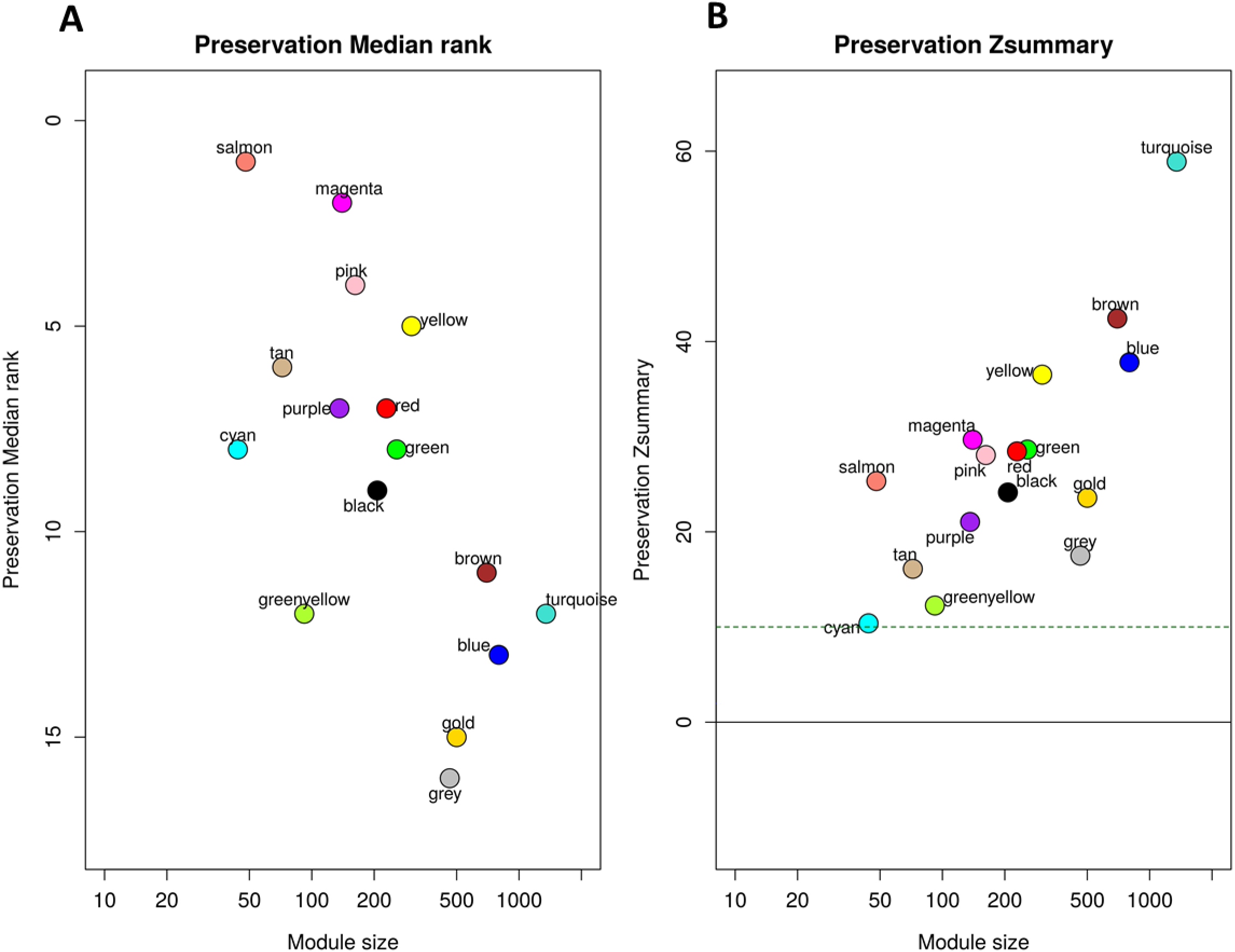
Preservation of discovery modules in validation cohort. **(A)** The median rank preservation statistics of the modules. Each module was represented by a point, labeled with the corresponding color and name. The Y-axis represents the median rank of observed preservation statistics per module, while the X-axis indicates the number of genes within each module. A low preservation median rank value indicates a high level of preservation. The gold module was an artificial module comprised of 500 randomly selected genes. The grey module consisted of non-connected genes identified in the WGCNA analysis of the discovery cohort. These two modules exhibited the highest median rank, indicating low preservation, which ensured our preservation analysis controlling well the background noise signal. **(B)** The Zsummary preservation statistics of the modules. Each module was represented by a point, labeled with the corresponding color and name. The Y-axis represents the Zsummary statistic of each module based on 1,000 permutations of module labels, while the X-axis indicates the number of genes within each module. The horizontal green dash line corresponds to the threshold of Zsummary>10 indicating strongly persevered modules. The presence of all modules in discovery were validated in validation cohort with Zsummary>10.

**Figure S3.**
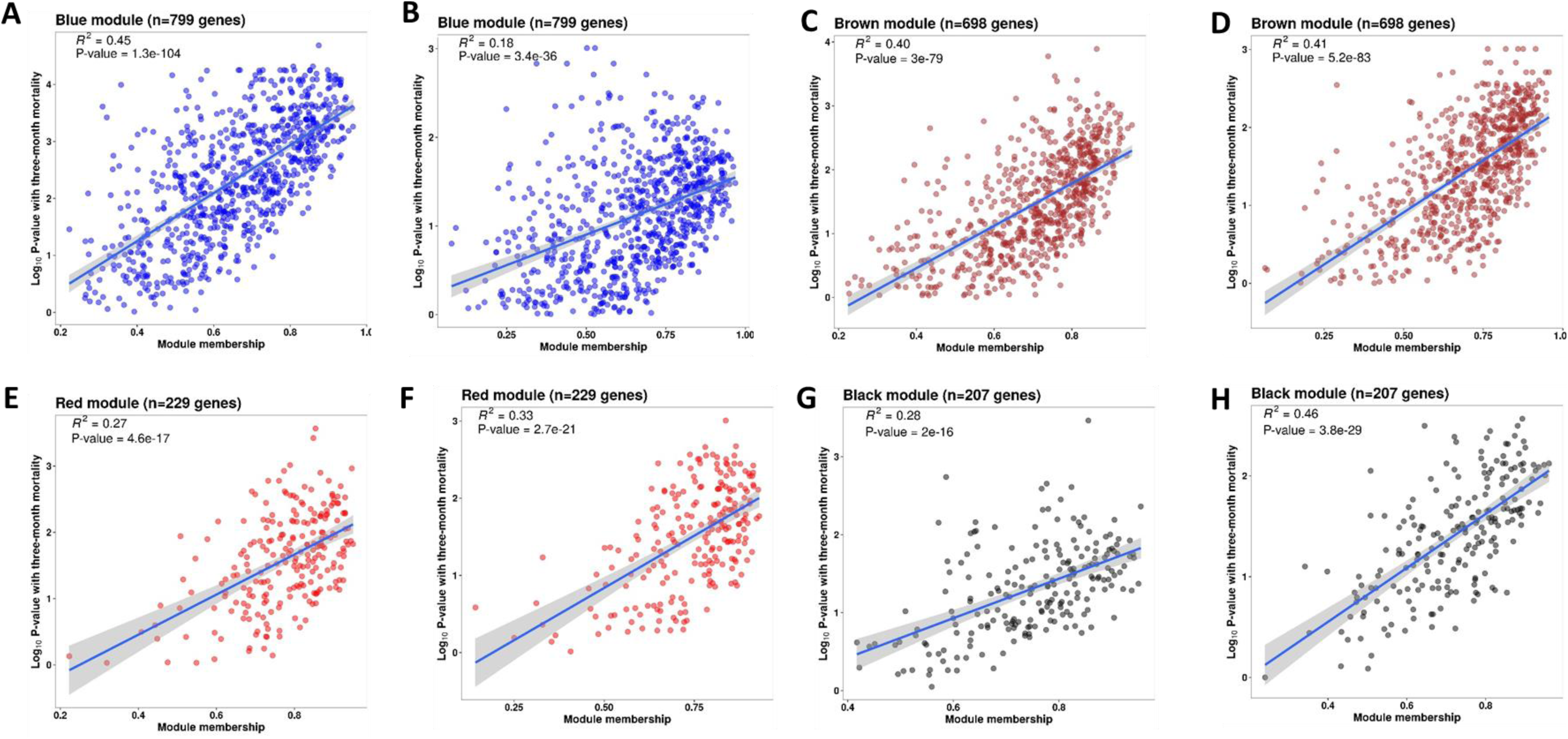
Correlations between gene module membership and gene significance with mortality in 4 associated modules in discovery and validation cohorts. **(A, C, E and G)** are scatter plots for blue, brown, red and black modules in discovery cohort. (**B, D, F and H)** are scatter plots for blue, brown, red and black modules in validation cohort. For each module, each dot represents one gene in the module. The X-axis represents module membership calculated by Pearson correlation between gene expression level and its corresponding PC1 of that module. The Y-axis represents –log_10_(p-value) of the association between gene expression and three-month mortality based on the Cox regression model adjusted for age, HIV, and dexamethasone treatment. Hub-genes were those on the right corner of the plot. The displayed R^2^ and P value were taken from linear regression models.

**Figure S4.**
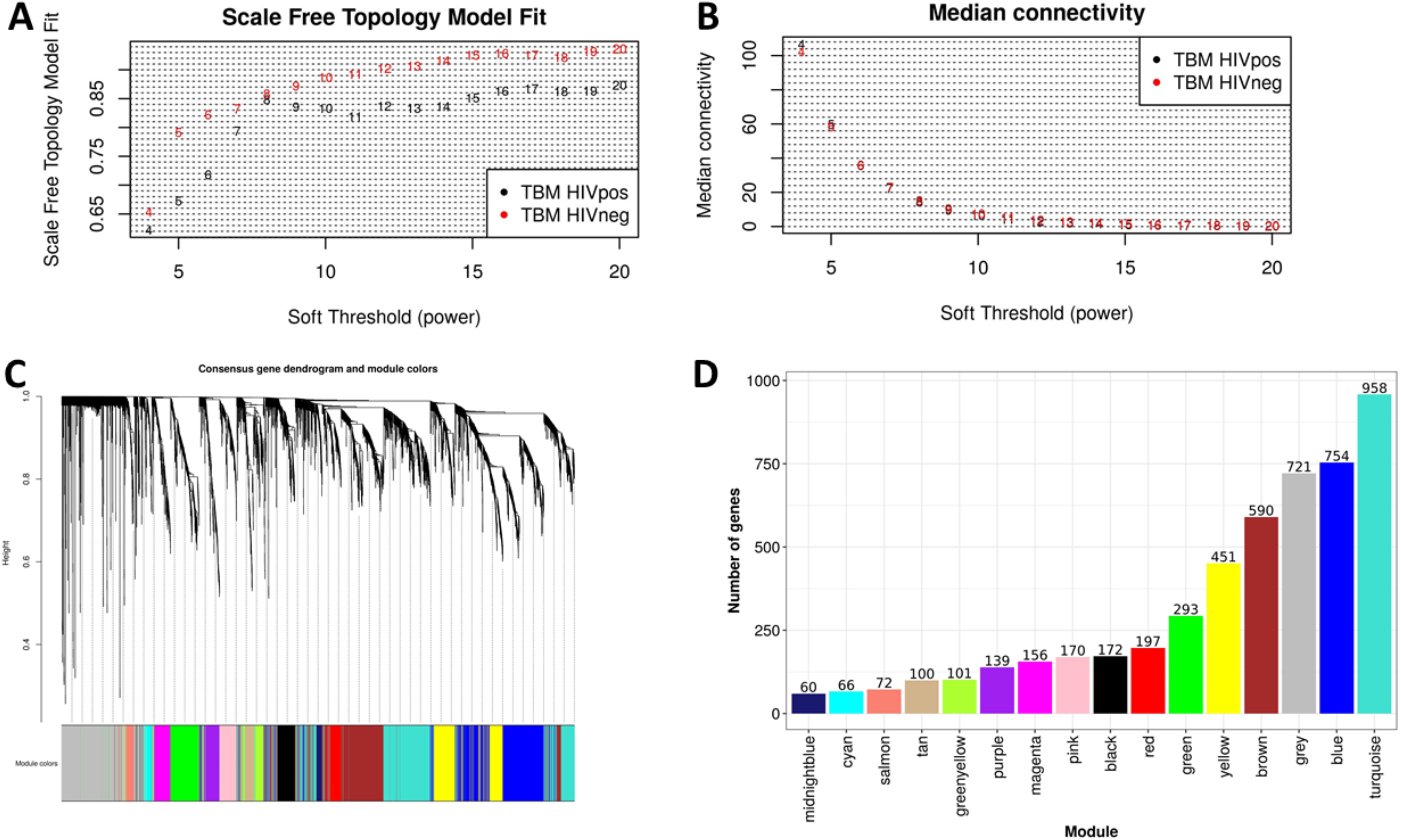
Construction of consensus WGCNA in HIV-negative (n=207) and HIV-positive (n=74) Consensus analysis of network topology for various soft-thresholding powers of the top 5,000 most variant genes based on the scale-free network model. **(A, B)** showed the summary network indices (y-axes) as functions of the soft thresholding power (x-axes). Numbers in the plots indicate the corresponding soft thresholding powers. The plots indicated that approximate scale-free topology is attained around the soft-thresholding power of 8 for both sets, which is the lowest power that satisfies the approximate scale-free topology criterion for both cohorts, HIV-positive (black dots) and HIV-negative (red dots). The adjacency matrix of the scale-free network between genes of each cohort was determined and scaled. The consensus topological overlap of two adjacency matrix were identified and served as dissimilarity metric for the hierarchical clustering. **(C)** displayed the dendrogram of the consensus gene co-expression network. Each cluster was referred as a module and assigned with a color. **(D)** the bar-plot indicates the number of genes contained in each module.

**Figure S5.**
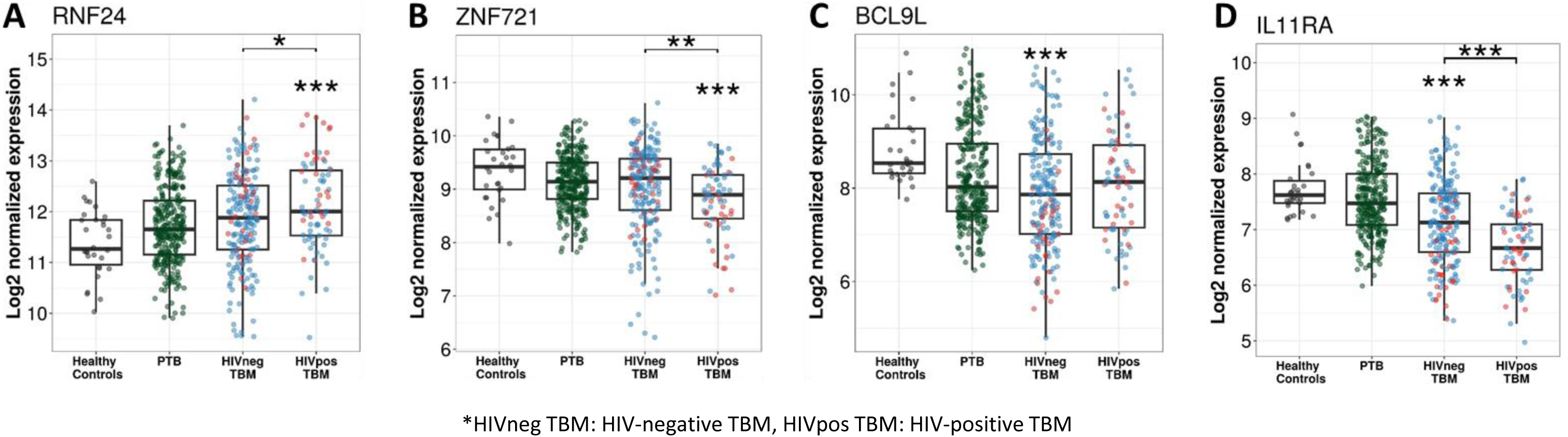
Gene expression of representative hub genes in healthy controls (n=30), PTB (n=295), HIV-negative TBM (n=207) and HIV-positive TBM (n=74) Each dot represents gene expression from one participant. **(A, B)** expression of RNF24 and ZNF721 hub genes from the specific blue module. **(C, D)** expression of BCL9L and IL11RA hub genes from the specific yellow modules. The box presents median, 25^th^ to 75^th^ percentile and the whiskers present the minimum to the maximum points in the data. Comparisons were made between dead (red) with survival (blue) or between HIV-negative and HIV-positive TBM by Wilcoxon rank sum test with p-values displayed as significance level above the boxes and the horizontal bars, respectively (* < .05, ** < .01, *** <.001).

**Figure S6.**
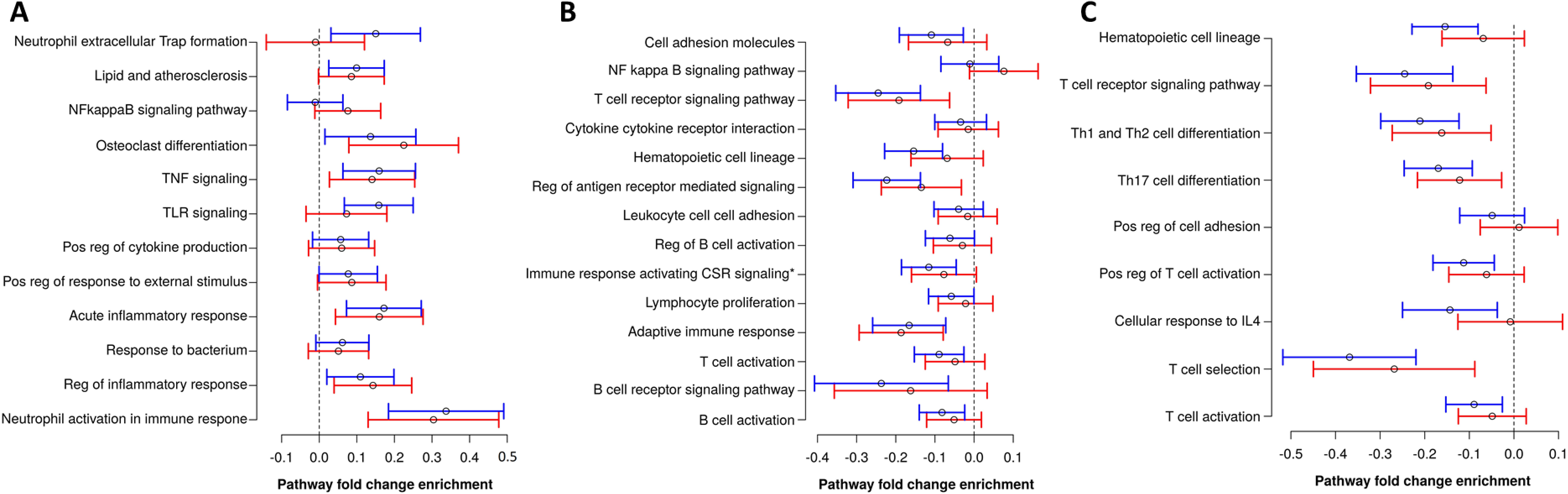
Pathway fold change enrichment to mortality of top hit pathways in blue, brown and red modules from the primary analysis. Pathway fold change enrichment to mortality of each biological pathway defined as the distribution of the total of mean differential expression between dead and survival TBM of all genes involved in the pathway based on Qusage method (Yaari G at el, 2013). Median and 95% confidence interval of pathway fold change of each top-hit biological pathway in the blue **(A)**, brown **(B)**, and red module **(C)** in the primary analysis. Blue lines are from HIV-negative cohort and red lines from HIV-positive cohort. Pathway fold change enrichment above 0 indicates up-regulation while below 0 indicates down-regulation in dead. Vertical dashed lines indicate no difference in dead compared to survival TBM.

**Figure S7.**
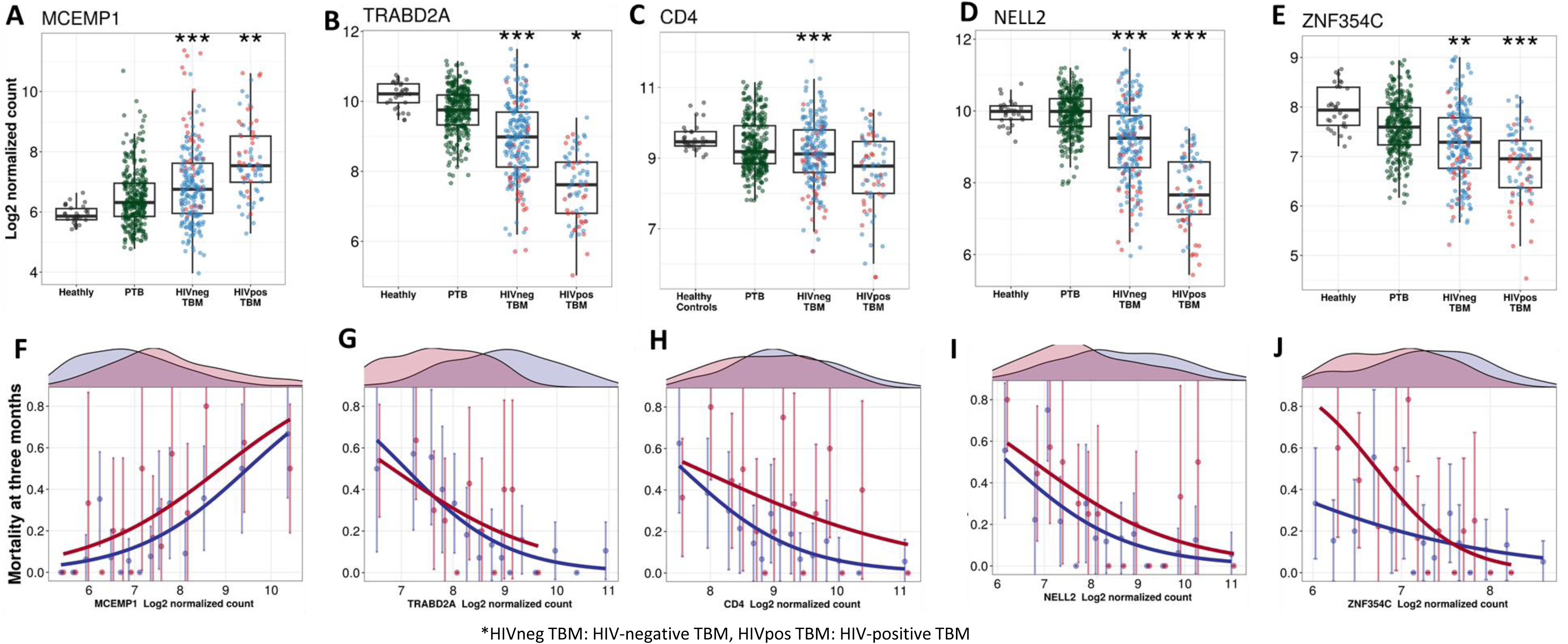
Gene expression and mortality of selected hub genes in TBM, PTB and healthy participants. **(A-E)** Boxplots visualized distribution of gene expression of MCEMP1, TRABD2A, CD4, NELL2 and ZNF354C in healthy (n=30), PTB (n=295), TBM without HIV (n=207) and with HIV (n=74). Boxes indicate median and inter-quantile range. Dots indicate data in individuals. The comparisons were made between dead (red) and survival (blue) by Wilcoxon rank sum test with p-value displayed as * < .05, ** < .01 and *** <.001. **(F-J)** Associations between mortality and gene expression of MCEMP1, TRABD2A, PKD1, NELL2 and ZNF354C. In each figure, the upper panel corresponds to gene distribution in HIV-negative (blue) and HIV-positive (red), the lower panel presents the approximation of the association between that gene and mortality. Their gene expression were divided into 15 groups using equal-distant quantiles (1/15,…, 14/15) intervals and the proportion of mortality within those groups of patients were computed. Each point (error bar) presented the proportion with its confidence interval of mortality per group. The line represents the logistic curve that illustrates the mortality trend corresponding to a two-fold increase in gene expression.

**Figure S8.**
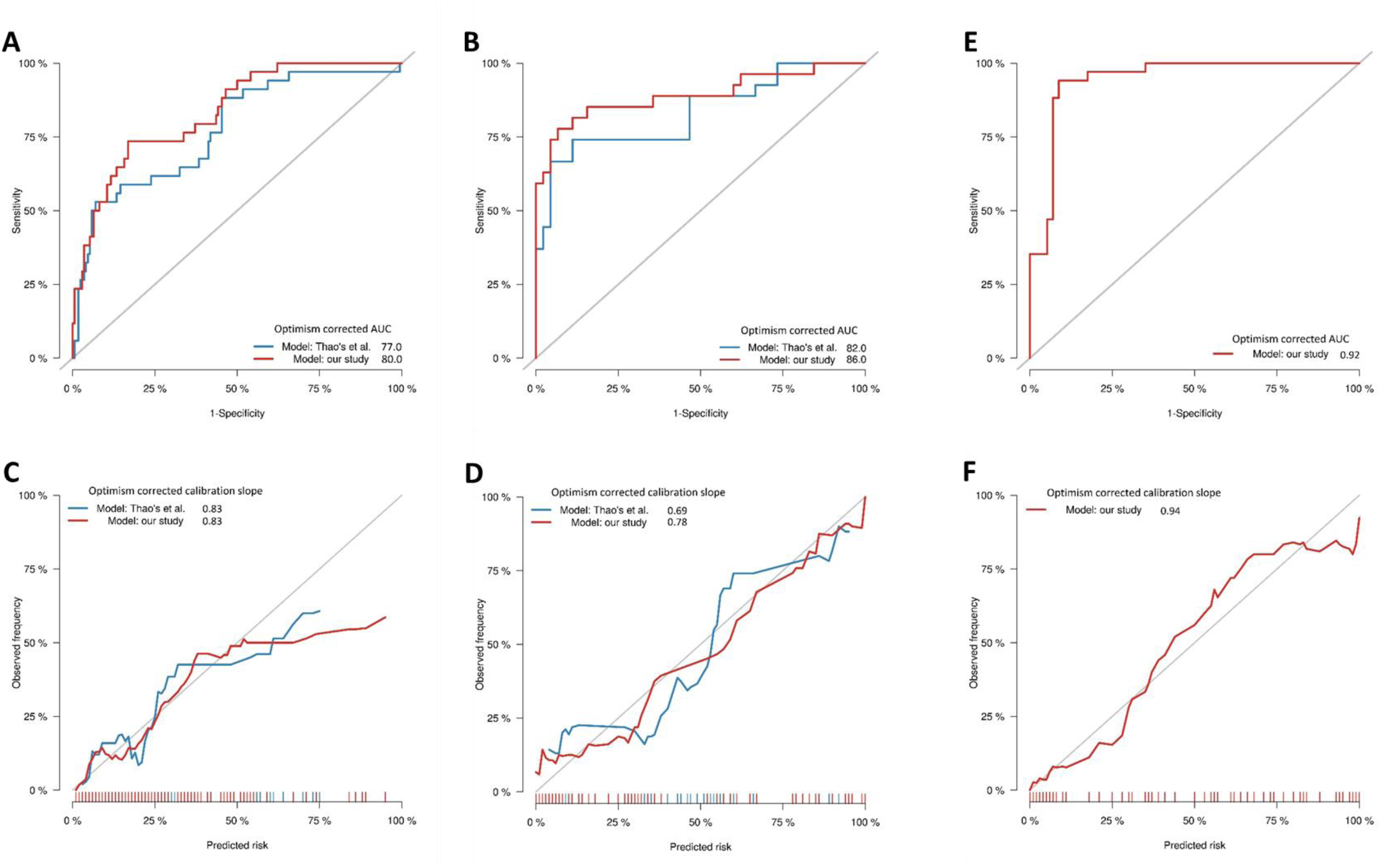
Performance of optimal gene set for TBM mortality prediction. Receiver operating characteristic (ROC) curves for three-month mortality for our developed model using expression level of 4 genes (MCEMP1, NELL2, CD4 and ZNF354C) with 2 clinical predictors (age, MRC grade) (red line) and the model develop by Thao et al. (PMID: 29029055) (blue line) for HIV-negative cohort **(A)**, for HIV-positive cohort **(B) and for HIV-negative PCR validation cohort (E)**. The optimism corrected area under the curve (AUC) was calculated based on 1,000 times bootstrap subsampling as described in the methods. Calibration plots were presented for the corresponding prediction models for HIV-negative **(C)**, HIV-positive cohort **(D) and HIV-negative PCR validation cohort (E)**.

**Figure S9.**
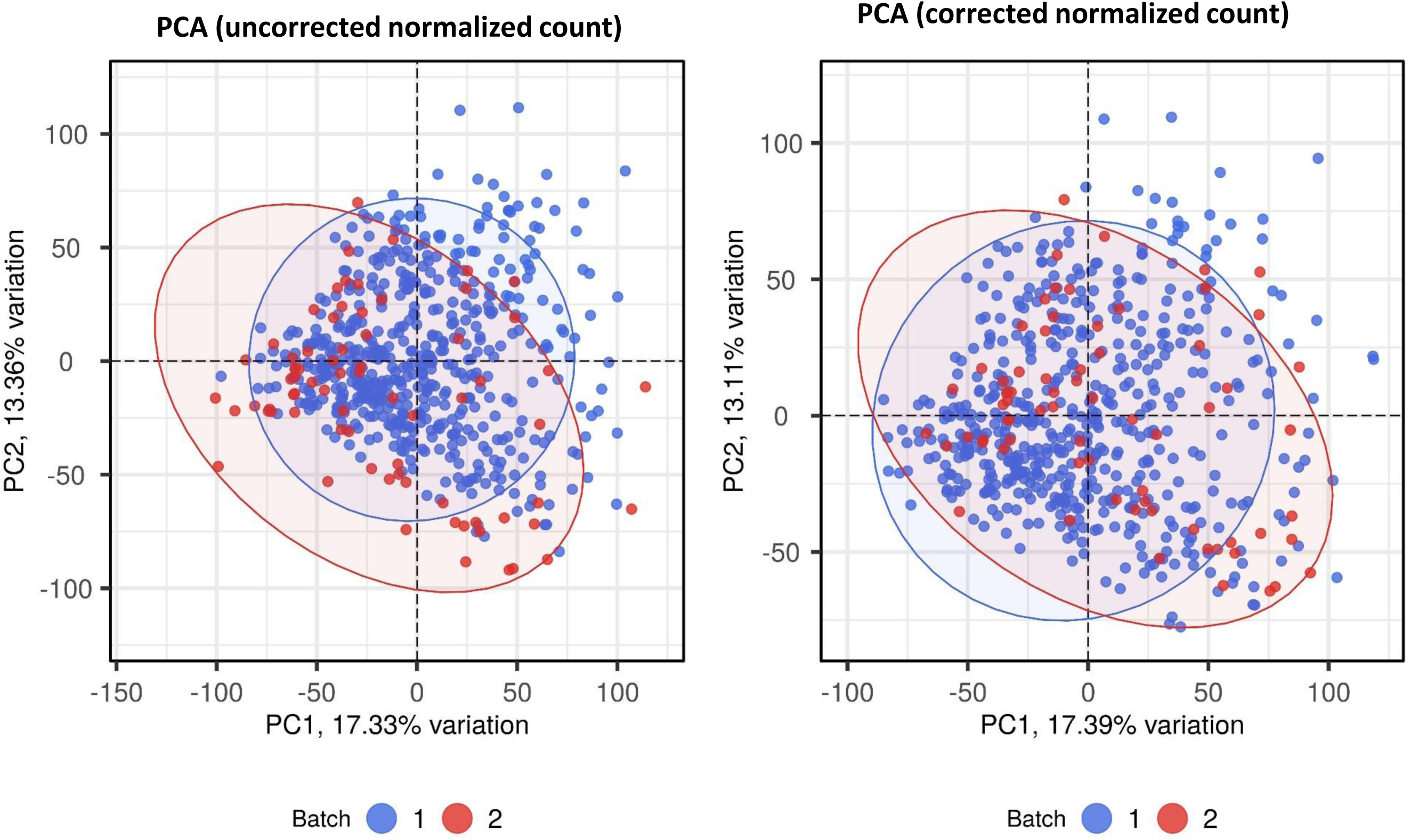
Batch correction for RNA-seq normalized count data. Principle component analysis (PCA) of transcriptomic data before (A) and after batch correction by combat function in SVA r package. Each symbol represents one individual with color coding different cohorts. The x-axis represents principle component (PC) 1, while y-axis represents PC2

**Figure S10.**
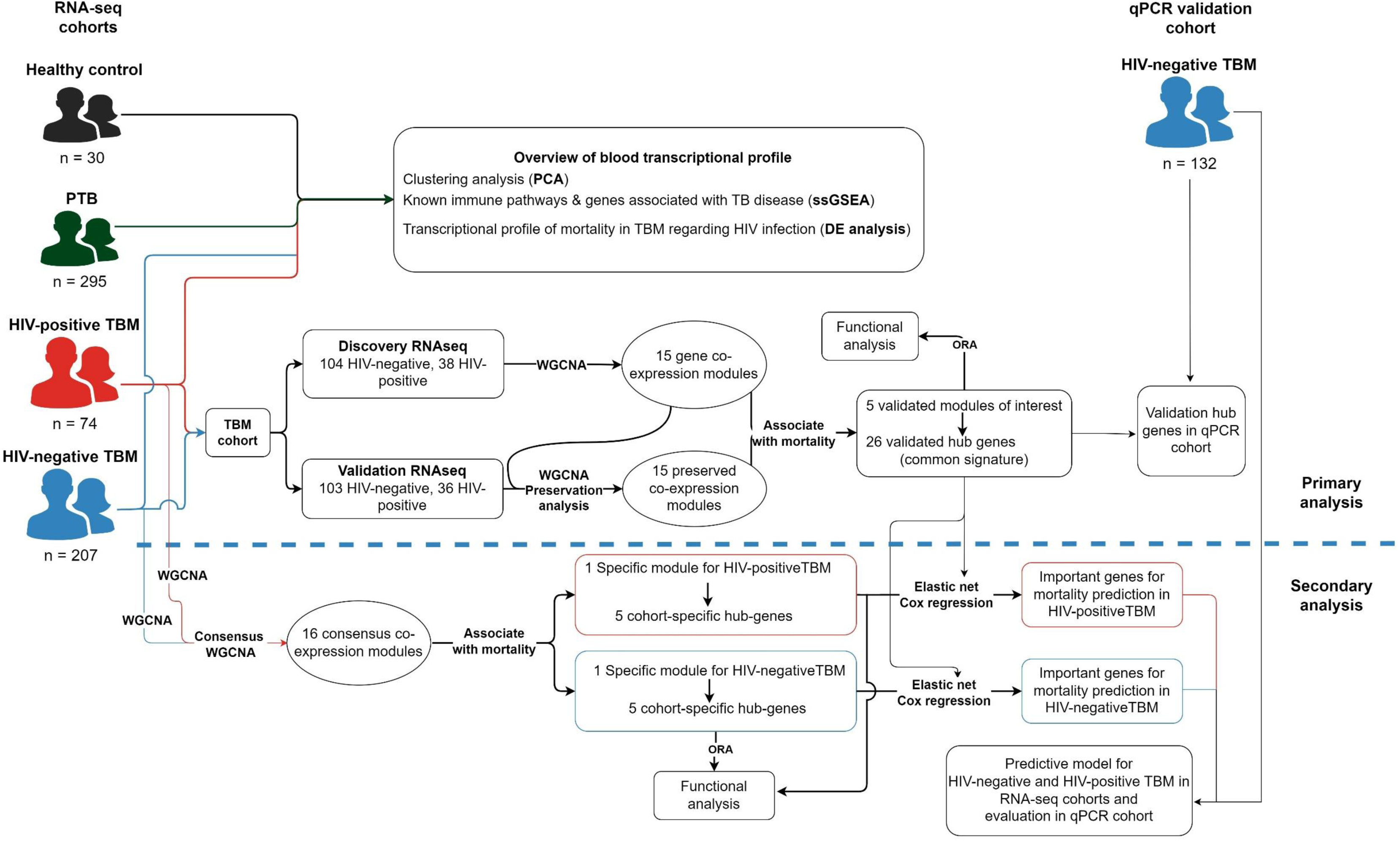
Analysis workflow diagram. TBM participants were recruited from two randomized control trials for adjunctive dexamethasone treatment of HIV-negative and HIV-positive adults with TBM (*LAST ACT: NCT03100786; ACT HIV: NCT03092817). Samples from 207 HIV-negative and 74 HIV-positive patients were used for RNA sequencing and randomly divided into discovery and validation cohorts to identify biological pathways and hub genes associated with three-month mortality.

